# The miR-430 locus with extreme promoter density is a transcription body organizer, which facilitates long range regulation in zygotic genome activation

**DOI:** 10.1101/2021.08.09.455629

**Authors:** Yavor Hadzhiev, Lucy Wheatley, Ledean Cooper, Federico Ansaloni, Celina Whalley, Zelin Chen, Stefano Gustincich, Remo Sanges, Shawn Burgess, Andrew Beggs, Ferenc Müller

**Author notes:** These authors contributed equally (Yavor Hadzhiev, Lucy Wheatley).

## Abstract

In anamniote embryos the major wave of zygotic genome activation (ZGA) starts during the mid-blastula transition. This major wave of ZGA is facilitated by several mechanisms, including dilution of repressive maternal factors and accumulation of activating transcription factors during the fast cell division cycles preceding the mid-blastula transition. However, a set of genes escape global genome repression and are activated substantially earlier, during what is called, the minor wave of genome activation. While the mechanisms underlying the major wave of genome activation have been studied extensively, the minor wave of genome activation is little understood. In zebrafish the earliest expressed RNA polymerase II (Pol II) transcribed genes are activated in a pair of large transcription bodies depleted of chromatin, abundant in elongating Pol II and nascent RNAs (Hadzhiev *et al.*, 2019; Hilbert *et al.*, 2021). This transcription body includes the miR-430 gene cluster required for maternal mRNA clearance. Here we explored the genomic, chromatin organisation and *cis-*regulatory mechanisms of the minor wave of genome activation occurring in the transcription body. By long read genome sequencing we identified a remarkable cluster of miR-430 genes with over 300 promoters and spanning 0.6 Mb, which represent the highest promoter density of the genome. We demonstrate that the miR-430 gene cluster is required for the formation of the transcription body and acts as a transcription organiser for minor wave activation of a set of *zinc finger* genes scattered on the same chromosome arm, which share promoter features with the miR-430 cluster. These promoter features are shared among minor wave genes overall and include the TATA-box and sharp transcription start site profile. Single copy miR-430 promoter transgene reporter experiments indicate the importance of promoter-autonomous mechanisms regulating escape from global repression of the early embryo. These results together suggest that formation of the transcription body in the early embryo is the result of high promoter density coupled to a minor wave-specific core promoter code for transcribing key minor wave ZGA genes, which are required for the overhaul of the transcriptome during early embryonic development.

## INTRODUCTION

The major wave of zygotic genome activation(ZGA) is characterized by activation of thousands of genes when the dividing cells reach a threshold nucleo-cytoplasmic ratio (Reviewed in (Lee *et al.*, 2014; Schulz and Harrison, 2019; Vastenhouw *et al.*, 2019)). This nucleo-cytoplasmic ratio has been suggested to reflect competition between the transcription machinery and maternally deposited repressing chromatin factors. These repressive factors include maternally-deposited excess of histones, which get diluted by exponential cell divisions (Amodeo *et al.*, 2015; Joseph *et al.*, 2017).}. In addition, several molecular mechanisms have been suggested to regulate the timing of genome activation: i) delayed availability of pioneering transcription factors (Lee *et al.*, 2013); ii) or that of the transcription initiation machinery (Bartfai *et al.*, 2004; Ferg *et al.*, 2007; Veenstra *et al.*, 1999); iii) sequential increase in chromatin accessibility (Blythe and Wieschaus, 2016; Palfy *et al.*, 2020); iv) gradual formation of transcriptionally competent chromatin (Lindeman *et al.*, 2011; Chan *et al.*, 2019); v) elongation of the cell cycle, permissive for elongating transcription (Blythe and Wieschaus, 2015; Collart *et al.*, 2013).

However, many genes escape the global repression and get activated in advance of the rest of the genome, during what is called the minor wave of genome activation. In zebrafish the minor wave of genome activation includes miR-430 (Abe *et al.*, 2018; Heyn *et al.*, 2014), a key regulator of maternal mRNA clearance in zebrafish (Giraldez *et al.*, 2006). It is currently unclear what mechanisms drive selective activation of these ‘minor wave’ genes and therefore it is important to characterize their epigenomic features, which may inform on various genome activation mechanisms involved. In zebrafish, the minor wave of genome activation starts at 64-cell stage and reaches high levels of activity by 256-512-cell stage, during the interphase of short semi-synchronous cell cycles and characterised by high activity of miR-430 primary transcript production (Chan *et al.*, 2019; Hadzhiev *et al.*, 2019). Notably, miR-430 nascent RNA (primary microRNA) accumulates in a ‘transcription body’, which carries most of the detectable, transcriptionally active, elongating Pol II and nascent RNA detectable during the minor wave of genome activation between 64- to 512-cell stages (Hadzhiev *et al.*, 2019). The high transcriptional activity of this body has been associated with local chromatin depletion (Hilbert *et al.*, 2021). The minor wave of gene activation includes *zinc finger* (*znf*) genes, the nascent RNAs of which colocalise in the transcription body with miR-430 RNAs and thus, this body represents an accessible model to address the shared and distinctive mechanisms of the activation of the very first genes in the embryo. We hypothesized that the genetic and epigenetic mechanisms underlying the activation of miR-430 locus may explain the unique formation of this transcription body. Additionally, the genomic locus of miR-430 genes is marked by CTCF and Rad21 (factors involved in chromatin looping and higher-order structure) (Meier *et al.*, 2018), suggesting chromosome conformation mechanisms involved in its activation. MiR-430 activation is dependent on BRD4 and competent chromatin marked by histone H3K27 acetylation (ac) (Chan *et al.*, 2019; Miao *et al.*, 2020) and on the pioneer transcription factors (also called stem cell factors): Nanog, Pou5f3, Sox19b (Chan *et al.*, 2019; Leichsenring *et al.*, 2013; Miao *et al.*, 2020). These stem cell factors have been shown to activate main wave of ZGA in fish (Lee *et al.*, 2013; Leichsenring *et al.*, 2013; Palfy *et al.*, 2020; Veil *et al.*, 2019) and homologs Pou5f3 and Sox3 in Xenopus (Gentsch *et al.*, 2019), while their role in the minor wave of genome activation is not yet understood.

Deposition of histone post-translational modifications (H3K4me1, H3K4me3, and histone variants (H2AZ) associated with gene regulation, precede genome activation (Lindeman *et al.*, 2011) and can even be detected in gametes (Wu *et al.*, 2011). These epigenetic dynamics suggest potential inheritance from the parents and raising the possibility of potentially instructive role for regulation of both the minor and main wave of genome activation (Liu *et al.*, 2018; Murphy *et al.*, 2018; Wike *et al.*, 2021; Zhang *et al.*, 2018; Zhu *et al.*, 2019)

In this study we have explored the molecular mechanisms underlying the activation of minor wave genes using the miR-430 gene locus and surrounding early acting genes. Here we demonstrate that the miR-430 gene cluster is required for the formation of the transcription body and acts as a transcription organiser for early wave activation of a set of *znf* genes, which share promoter features with the miR-430 cluster, however, reside outside of the cluster on the same chromosome arm. In addition, we show that minor ZGA-specific core promoters with distinct architecture from that used by the major wave of ZGA, together with high promoter density, are necessary for the formation of a transcription body, to transcribe minor ZGA genes, required for correct embryonic development. While promoter density enables the formation of a transcription body, single copy genes with promoters, which share features with those in the transcription body can also escape global transcriptional repression and are able to become activated in the minor wave.

## RESULTS

### Long read sequencing mediated genome assembly reveals the highest promoter density gene cluster in the zebrafish genome

In the current genome assembly (GRCz11) the miR-430 locus is a multicopy gene cluster, which consist of 8 complete miR-430 primary transcript gene units, each containing a 650 bp promoter region and triplets of precursor miRNAs, which themselves are repeated in 2 (duplex) or 3 times (triplex) per promoter (Hadzhiev *et al.*, 2019) (**Figure 1A**). Each precursor miRNA triplet consists of 4 different subtypes of precursor genes: a, b, c and the ‘a’ variant (called ‘i’), organised into repetitive units of 3 precursors (Hadzhiev *et al.*, 2019). In the reference genome assembly (GRCz11), the 5’end of the locus is truncated, missing the promoter sequence, which suggests that this section of the assembly is incomplete, probably due to the difficulty of assembling the highly repetitive structure (**Figure 1A**). Therefore, we embarked to reassemble the the miR-430 locus and we generated Oxford Nanopore long read sequencing data from AB male zebrafish at ~44x coverage and complemented this with long read sequencing using PacBio (Pacific Bioscience) sequencing from a heat shock diploid “double heterozygous” (Westerfield and ZFIN., 2000) Tuebingen-AB hybrid female zebrafish at ~78x coverage.

**Figure 1.**
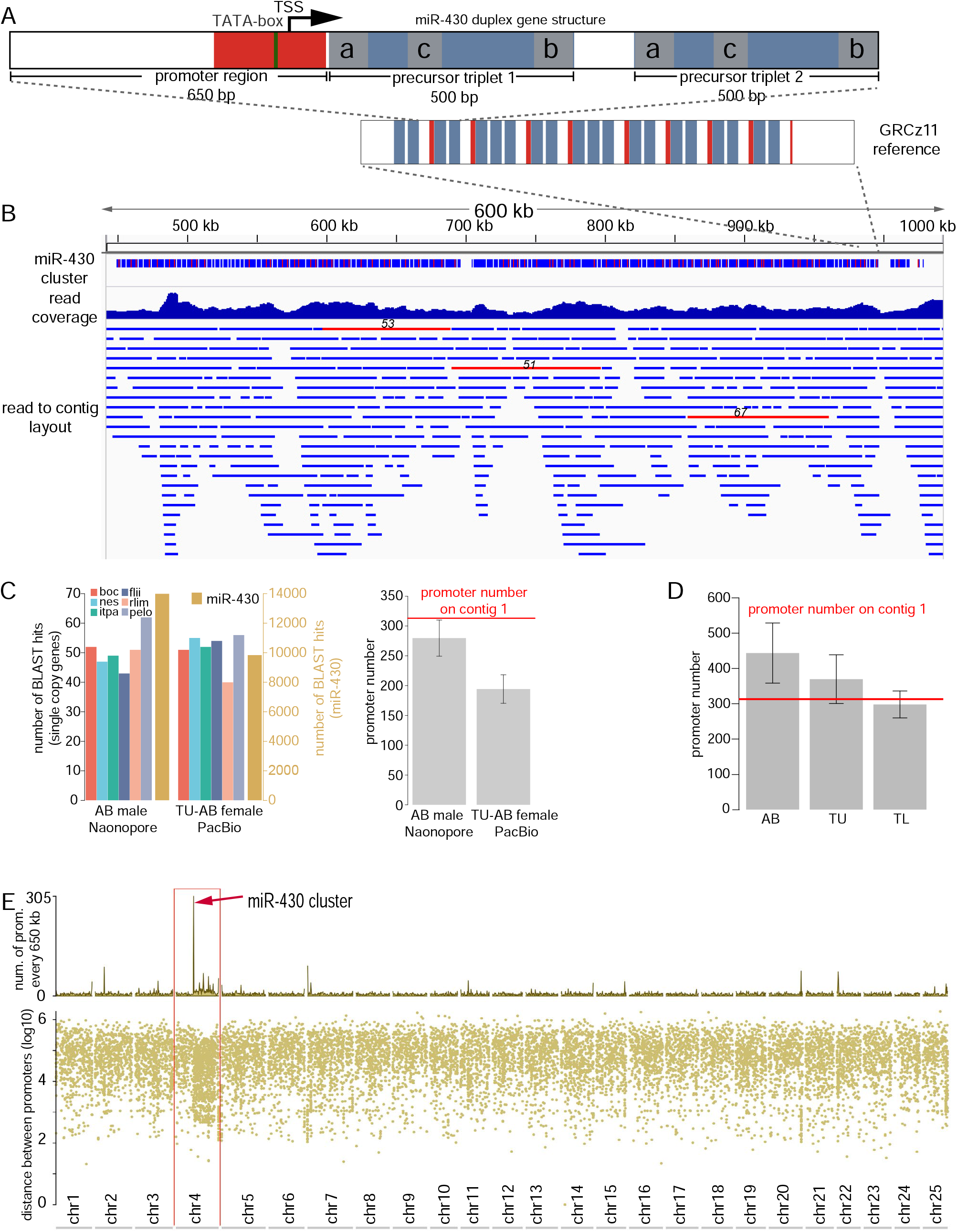
De novo assembly of the miR-430 cluster from long read sequencing data. **A,** Schematic representation of the most common miR-430 gene structure (top panel), the duplex consisting of promoter region (core promoter marked in red) and two pre-miRNA triplets (blue) each containing 3 of the of pre-miR-430 subtypes: miR-430a, miR-430b and miR-430c (gray). Bottom panel: the miR-430 gene cluster structure in current reference assembly (GRCz11). Promoter and triplet structures are represented by red and blue boxes, respectively. **B**, Genome Browser screenshots of the miR-430 gene cluster assembled from Nanopore long reads. The three tracks show a low resolution continuity structure of the gene cluster (top), long read coverage histogram (middle) and the read to contig assembly layout (bottom). Raw reads with 50 or more promoters are shown as red horizontal bars with the promoter number indicated on top of them. **C,** Left panel: number of BLAST hits in raw long reads of single gene promoters (right axis, black) and miR-430 (left axis, orange). Right panel: Estimated miR-430 promoter number normalised to that of single copy genes. The red line indicates the promoter number on the assembled contig on panel C and D. Error bars are standard deviation from the mean of the miR-430 promoter estimate normalised to each of the 6 genes. Gene symbols are indicated in distinct colours and represent the bar colours. **D** Estimated miR-430 promoter number by qPCR(absolute quantification) for three zebrafish strains. Error bars are standard deviation from the mean from three technical replicates. **E,** Rainfall plot representing the promoter density across each chromosome of the newly assembled zebrafish genome (GRCz11 reference guided). Red arrow indicates the mirR-430 gene cluster; Abbreviations: TSS, Transcription Start Site; AB, TU-AB, TU, TL are zebrafish strains used in C and D.

Initial analyses of the raw Nanopore long read data revealed 7 raw reads, which contained 50 or more miR-430 promoters (red highlight **Figure 1B** and **Supplementary Figure 1A**), with 74 promoters on a single 111kb long raw read with 128 triplet precursors. The longest read spanning the miR-430 cluster was ~122 kb long with 67 promoters and 149 triplets. In the PacBio data the read with the highest promoter number was ~23.5 kb long and contained 13 promoters and 26 triplets. The longest read spanning the locus was ~34 kb long and contained 10 promoters and 39 triplets.

Next a *de novo* genome assembly was generated using the Canu assembler (Koren *et al.*, 2017). Assembly of the Oxford Nanopore reads resulted in 2 contigs containing the miR-430 gene cluster, likely representing allelic variants. The first contig was ~3.44 Mb long with 313 miR-430 promoters and 655 triplet structures spanning ~577 kb (**Figure 1B**). The second contig was ~811 kb long containing 306 miR-430 promoters and 587 triplet structures spanning ~534 kb (**Supplementary Figure 1A**). As demonstrated by comparison of the read to contig layouts, of the two contigs presented in **Figure 1B** and **Supplementary Figure 1A**, contig 1 had overall better-read/coverage support and was used in further analyses.

The striking increase of copy number from 8 to over 300 repeated gene unit copies prompted us to seek independent validation of the copy number of the miR-430 primary transcript genes. To this end we estimated read coverage and compared to known single copy genes on the newly assembled genome. A BLAST search of the miR-430 promoter region (**Figure 1B**) and the promoter regions of 6 single copy genes as reference against the raw reads of our Nanopore and PacBio sequencing runs. An estimate of promoter number of miR-430 was generated by normalising the BLAST hit number to that of the single copy genes. This analysis led to a calculated copy number of 280±30 and 194±24 for each sequencing run respectively (**Figure 1C**). In addition, we estimated the miR-430 promoter number by qPCR using standard curve absolute quantification from three zebrafish stains (AB, TU and TL). The copy number estimated by this approach was 444±85, 370±69 and 298±38 for AB, TU and TL stains respectively (**Figure 1D**). The assembly of the PacBio reads did not result in continuous, end to end assembly of the miR-430 locus but was split between several (14) contigs (**Supplementary Figure 1B**). The total number of miR-430 promoters and triplet structures on all contigs from the PacBio data was estimated between 359 and 768.

While the copy number figures from the above approaches differ, but they are in a comparable range to that obtained by copy number calculation on the assembled contigs (**Figure 1C,D**) and taken together, indicate that the number of promoters present at the miR-430 locus is almost 2 orders of magnitude higher than that was mapped on the reference genome.

This large promoter number repeated every 1.8 kb on average in a ~0.6Mb region is an unusual feature and we sought to explore how this promoter density compares to the rest of the genome. To analyse promoter density in the zebrafish genome in general, we performed GRCz11 reference guided chromosome scaffolding of the haplotig-purged (removing one of the duplicates, allelic variant, conting, see Methods for details*) de novo* assembly from the Nanopore sequencing run. Promoters were identified by multimapping CAGE-seq data (Nepal *et al.*, 2013) to the newly scaffolded genome and promoter density was calculated. As demonstrated on **Figure 1E** the miR-430 cluster indeed shows by far the highest promoter density in the assembled genome, more than 4 times higher than other promoter dense regions and higher than any region of the compact genome of tetraodon pufferfish (data not shown). Intriguingly, promoter density is even generally higher than the rest of the genome on the chr4 long arm, which is known to carry multicopy genes (Howe *et al.*, 2016) and has unique gene density and chromatin features (White *et al.*, 2017). Taken together, long read sequencing-based genome assembly and read copy number estimation reveals almost two orders of magnitude higher copy number for the miR-430 gene locus than previously appreciated and provide a potential explanation for the timing and the scale of gene expression during the minor wave of genome activation, which appears in a large transcription body.

### Distinct epigenetic regulation of the miR-430 cluster and minor wave genes of genome activation

The occurrence of minor wave gene activation raises the question of what chromatin and DNA sequence features are contributing to its formation. To address this question, we first aimed to characterise the epigenetic profiles and promoter architecture features of the most highly expressed gene cluster at the minor wave of genome activation, miR-430 together in comparison to other genes, which are activated at distinct phases of developmental transcription programme. Publicly available epigenomic datasets representing chromatin opening (ATAC-seq) and associated with cis regulatory element regulation (H3K4me3-, H3K27me3- and H3K27ac-ChIP-seq) (Liu *et al.*, 2018; Zhang *et al.*, 2018; Zhu *et al.*, 2019), covering key early developmental stages were selected for analysis (see Methods for details).

The selected datasets were mapped to a custom genome, created by appending the miR-430 cluster containing scaffold from the Nanopore long-read assembly to the current zebrafish genome reference (GRCz11/danRer11) sequence file. To investigate the role of epigenomic features in the early genome activation, their signal levels and dynamics during early development (blastula stages) were compared among 5 distinct gene sets (25 genes each) selected based on the timing of their onset of activation (see Figure 2): 1) **MiR-430** genes: The earliest and most highly transcribed gene cluster during the first (minor) wave of zygotic genome activation. **2) Minor Wave (MiW)** genes: detected as activate during the first wave of the zygotic genome activation by Ethynyl Uridine (EU) incorporation RNA-seq (Heyn *et al.*, 2014).

**Figure 2.**
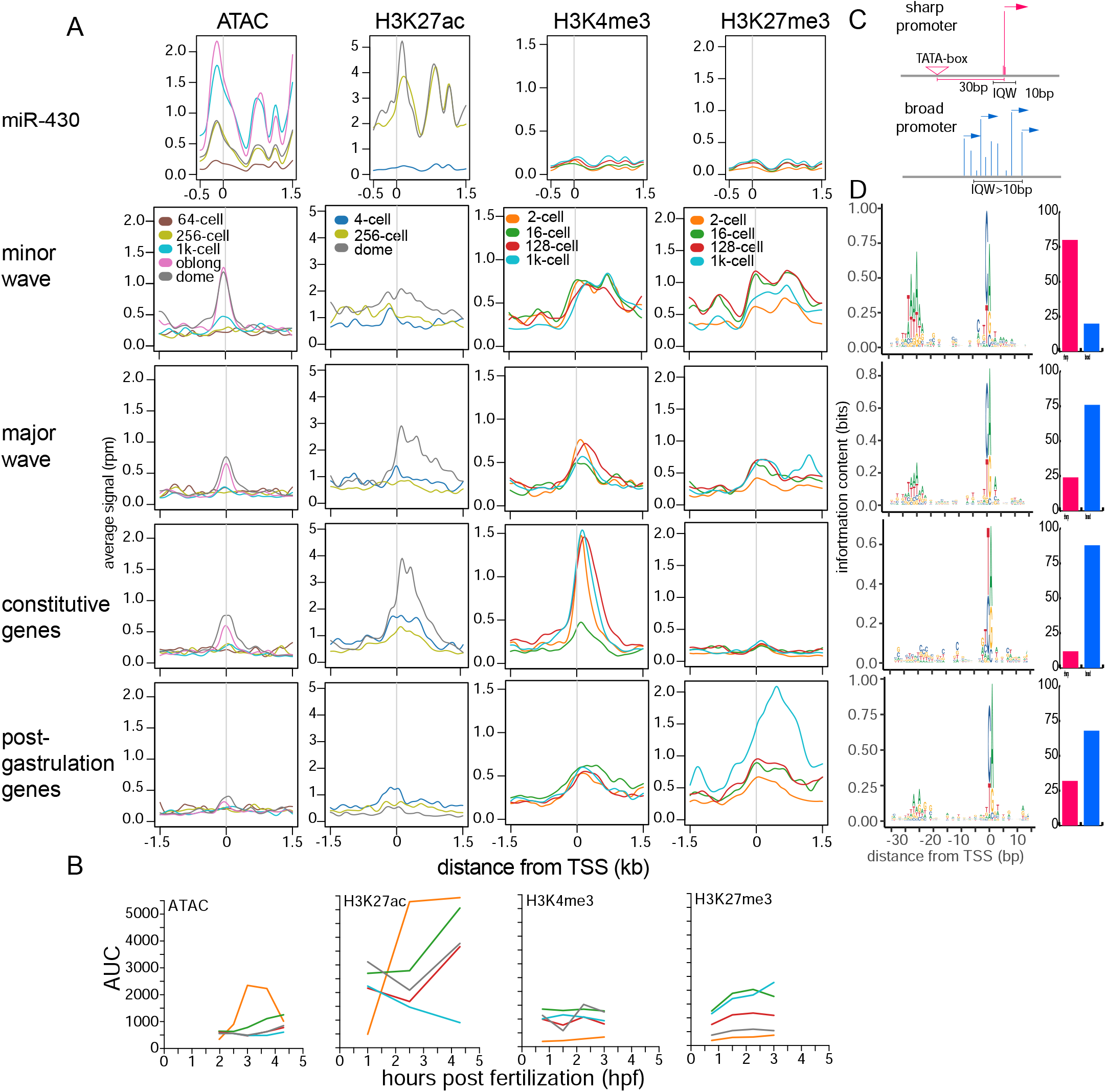
Epigenomic features of miR-430 cluster, compared to other gene sets activated at distinct phases of embryo development. **A,** Aggregation plots showing the signal distribution (mean) around the TSS (position 0) for each gene set (right side labels) and each epigenetic feature (top labels). **B,** Line graphs show the total signal (calculated as area under the curve from the aggregation plots on panel A) of each epigenetic feature for each gene set. **C,** Schematic comparing the sharp and broad promoter architectures. **D,** Promoter architecture comparison between the 4 gene sets. Left panels show sequence logos of the −35 +15 region relative to the TSS, right panel shows percentage of the genes with sharp (red) and broad (blue) promoter type in each gene set, determined by the interquartile width of the corresponding CAGE consensus cluster. Abbreviations: TSS: Transcription Start Site, MiW: minor wave, MjW: major wave, CG: constitutive genes, PG: post gastrulation set of genes, AUC: area under curve (used as a measure of total signal), IQW: interquartile width

Genes were selected with no maternal contribution and > 2.5 fold change between 128- and 512-cells. In many cases the pre-MBT expression could not be detected at significant levels by conventional RNA-seq and CAGE-seq. 3) **Major Wave (MjW)** genes: randomly selected genes activated at the major wave of ZGA and not detected during the minor wave (Heyn *et al.*, 2014). 4) **Constitutive Genes (CG):** housekeeping genes with maternal contribution and expressed throughout development. These genes were shown to use two different promoter codes present on the same core promoter platform (Haberle *et al.*, 2014). Firstly, a TATA-like motif called W-box was utilised in the oocyte, then later replaced by a nucleosome positioning signal of AT/GC enrichment boundary downstream of the transcription initiation start site (TSS) at ZGA. 5) **Post-gastrulation** genes **(PG):** activated after gastrulation during early segmentation stages. Expression dynamics for all gene sets were determined based on the CAGE-seq dataset (Nepal *et al.*, 2013) mapped to GRCz11/ danRer11, ENSEMBL v95 gene annotations (**Supplementary Figure 2 A-D**).

Next, the state (open/closed) of chromatin at the cluster was examined from the ATAC-seq datasets (**Figure 2 A,B).** The distribution profile of the signal was as expected for all datasets peaking 100-150 bp upstream of the TSS in the nucleosome free region of all active promoters. The signal levels on the miR-430 genes, as well as on the early and late zygotic gene sets, correlated with the gene expression dynamics, e.g., miR-430 is highly active early and shows the most open chromatin, whereas the minor wave genes were generally more open and become open at earlier developmental stages than the major wave genes. The major wave genes were comparable to the constitutive gene set in line with their onset of zygotic expression at and post-MBT. Next we compared the open/closed chromatin states with the appearances of promoter associated histone modification marks. The levels of the gene activity-associated H3K27ac mark (Creyghton *et al.*, 2010; Ernst *et al.*, 2011; Rada-Iglesias *et al.*, 2011) correlated with that of the open chromatin states observed (compare the H3K27ac signal to ATAC signal on **Figure 2A,B**). The miR-430 gene clusters showed the highest signal, already at 256-cell stage (**Figure 2A,B**). The minor wave gene set also showed higher levels and broader signal distribution than the major wave genes, which in contrast to the minor wave set, were more comparable to the constitutive genes. Like the ATAC-seq profile, the lowest levels were observed in the post gastrulation set, in line with being inactive at these stages.

Despite the high degree of open chromatin and enrichment in H3K27ac corresponding to the high expression levels at early blastula stages, the miR-430 cluster showed significantly lower levels of the active or poised promoter mark H3K4me3 (Haberle *et al.*, 2014; Heintzman *et al.*, 2007; Lindeman *et al.*, 2011; Vastenhouw *et al.*, 2010) in comparison to all other gene sets. Although, when signal levels were compared internally between stages, the signal levels correlated with miR-430 activity (**Figure 2 A,B**). The highest levels H3K4me3 were observed in the constitutive and minor wave gene sets, with the latter also having a broader distribution of signal (**Figure 2 A,B**) in line with their activity states. The major wave genes showed lower signal at 256-cell stage, in line with their later expression initiation at sphere/ dome stages. Although not expressed at early blastula and only becoming active at significantly later (early segmentation) stages, the post gastrulation genes showed relatively high H3K4me3 signal, comparable to the major wave genes (**Figure 2 A,B**).

An interesting dynamic of H3K4me3 signal was also observed between developmental stages. Relatively high levels were observed at the 2-cell stage with a significant drop at 16-cell, most prominent in the constitutive and major wave gene sets. This dynamic indicates potential parental inheritance or early embryonic pre-marking, that is followed by subsequent erasure and re-deposition at later stages, as previously reported (Zhu *et al.*, 2019). However, this “reprogramming” was not observed in the minor wave and post-gastrulation gene sets.

In summary, minor wave genes show broadly similar epigenetic profiles to the major wave genes, but with higher H3K27me3 levels. However, there is a notable difference between miR-430 and other minor wave genes in the apparent lack of H3K4me3 at this locus.

Previously, it was shown that zygotic genome activation in *Drosophila* is characterised by enrichment for the TATA-box (Chen *et al.*, 2013) and often associated with sharp or focussed transcription initiation profiles. In further pursuing potential distinctive features of minor wave genes, which may explain their distinct activation profiles, we analysed core promoter architecture features (**Figure 2C)** of minor wave genes of genome activation in relation to the 3 other gene activation groups. Sequence analysis of the 4 gene groups indicated marked enrichment for TATA-box at the canonical distance from the main TSS and corresponding sharp TSS architecture in minor wave genes, distinct from the major wave genes characterised by lack of canonical TATA-box and broader initiations site usage **(Figure 2D**)

This result together suggests that miR-430 promoters carry shared features with minor wave genes such as sharp promoters with TATA-box as well as distinct promoter features such as the lack of H3K4me3 signals. Both sets of minor wave genes are distinguished from the major wave genes by the TSS-determining TATA-box or TATA-like sequence signals and sharp TSS distribution.

### The proximal promoter sequences of the miR-430 gene cluster contain regulatory information sufficient to activate early transcription in a single copy and in chromosomal environment-independent manner

To identify the promoter autonomous sequence features that may inform on the regulation of miR-430 gene expression we have remapped the binding data of transcription factors previously described as regulators of the major wave of zebrafish genome activation on our new genome assembly (Lee *et al.*, 2013; Leichsenring *et al.*, 2013; Miao *et al.*, 2020). Publicly available ChIP-seq datasets (Leichsenring *et al.*, 2013; Xu *et al.*, 2012) for these stem cell factors were mapped on the de novo assembled miR-430 contig and signal distribution analysed. As demonstrated on Figure 3A the majority of the Nanog signal was distributed over the promoter region, upstream of the TSS as determined by CAGE-seq. Pou5f3 and Sox19b showed broader signal distribution (Sox19b in particular), but with main peaks over the promoter region. The region up- and downstream of miR-430 cluster show significantly less signal compared to the miR-430 promoter regions, suggesting that these pluripotency factors mainly bind proximally, with no evidence of distal enhancers **(Figure 3A and Supplementary Figure 3A**). In addition, we performed in silico transcription factor binding site analysis using the ClusterBuster software which identifies clusters of pre-specified motifs in a DNA sequence (Frith *et al.*, 2003). ClusterBuster analysis with the Nanog, Pou5f1::Sox2 and Sox2 motifs on the miR-430 contig resulted in the miR-430 annotated region being identified as one single cluster, enriched for high scoring binding sites (**Figure 3A** and **Supplementary Figure 3A**). The predicted binding sites for Nanog and Pou5f1::Sox2 were predominantly located in vicinity of the TSSs, with a Nanog motif at the peak of Nanog ChIP-seq signal (**Figure 3A**) and a high scoring Pou5f1::Sox2 site at ~300 bp upstream of the TSS (**Figure 3A**).

**Figure 3.**
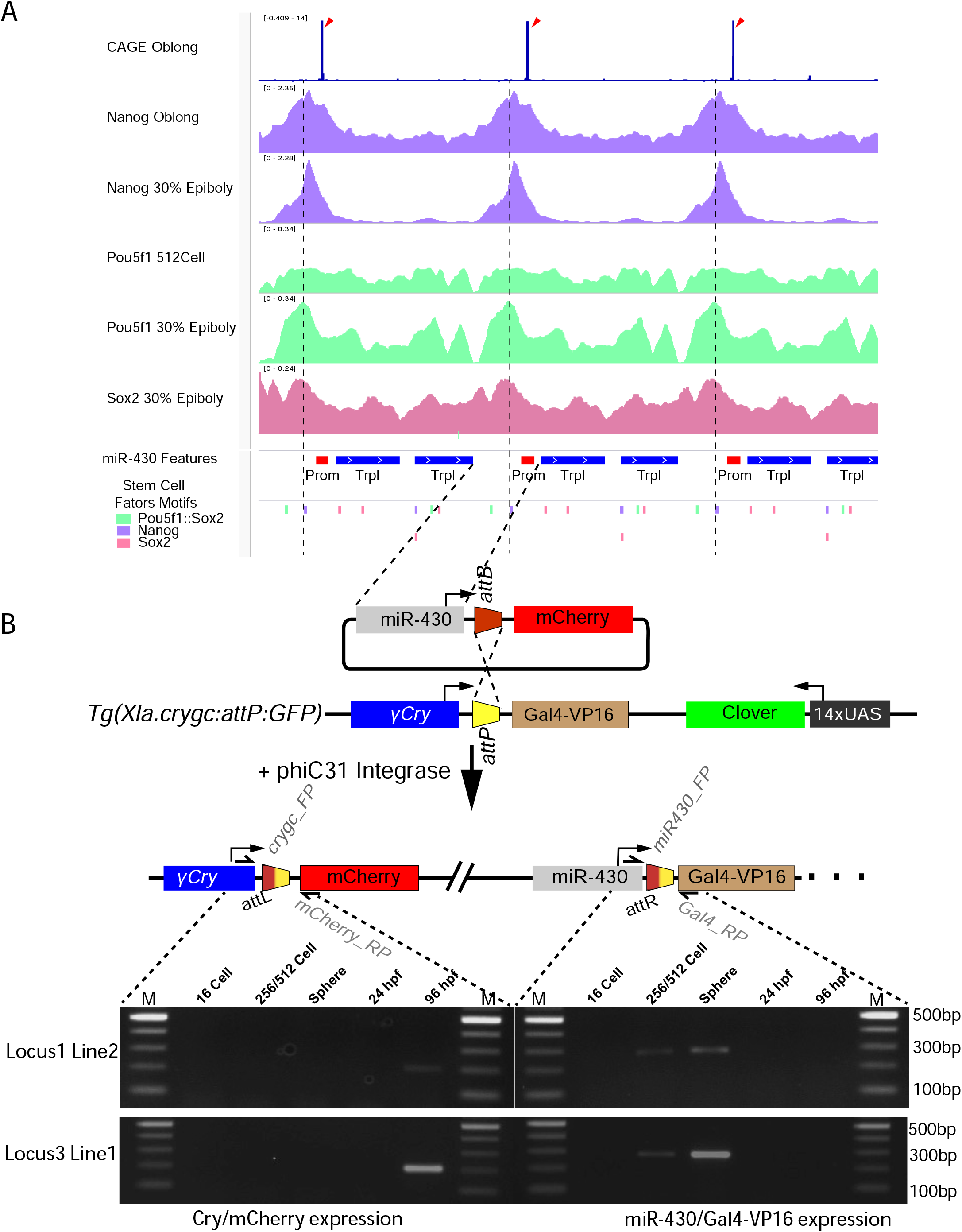
A single copy of the miR-430 promoter including stem cell transcription factor binding sites is sufficient to activate reporter expression during the minor wave of genome activation. **A,** browser screenshot showing CAGE-seq signal (top track, blue bar) marking the position of the TSS (red arrowheads).ChIP-seq signal coverage of the pluripotency factors Nanog (purple), Pou5f1 (green) and Sox2 (pink) are shown below. Bottom annotation tracks show the miR-430 gene features, with the core promoter marked in red and the precursor triplet in blue and the position of pluripotency factor binding sites, predicted by Cluster Buster. Purple dash line marks the Nanog binding site in the core promoter region close to the peak of the Nanog ChIP-seq signal. **B,** Top panel represents a schematic of the generation of miR-430 promoter-containing stable transgenics reporter lines using the PhiC31 site specific integration system. Bottom panel shows expression of the reporter miR-430 promoter (left) and γ-crystalline promoter used as control (right), detected by RT-PCR at the indicated stages. Abbreviations: TSS, Transcription Start Site, γ-Cry-(γ-crystalline C)

The highly multicopy nature of the miR-430 locus may provide high density of *cis-*regulatory elements locally to counter the repressive effect of maternally deposited negative regulators of transcription, such as abundance of maternal histones (Amodeo *et al.*, 2015; Joseph *et al.*, 2017). To test if the multicopy state is required or the sequence information of a single miR-430 promoter is autonomously sufficient to drive activity at the minor wave of genome activation we designed an experiment to integrate a single copy of the miR-430 promoter with a reporter gene into a heterologous genomic location. A 650 bp proximal promoter fragment extending in both directions to the upstream and downstream triplet structures from the CAGE-seq determined TSS, respectively, was isolated for functional analysis. This fragment contains the TATA-box and binding sites of Nanog and Pou5f1::Sox2 detected by ChiP-seq signal and predicted in silico (**Figure 3B**). This promoter sequence was cloned into a reporter plasmid used to generate stable transgenic lines by utilising the phiC31 targeted transgene integration system (Hadzhiev *et al.*, 2016; Roberts *et al.*, 2014) (**Figure 3B**). We generated several docking lines *Tg(Xla.crygc:attP--Gal4vp16,14UAS:Clover)UoBL1* and *Tg(Xla.crygc:attP--Gal4vp16,14UAS:Clover)UoBL3* that were used as different chromosomal landing sites. The integration site of the *Tg(Xla.crygc:attP--Gal4vp16,14UAS:Clover)UoBL3* line is on chr17 ~7 kb upstream of *slc16a9a* and ~6kb downstream of *npy4r (*chr17:20640317-20646440, danRer11)and is undetermined for the *Tg(Xla.crygc:attP--Gal4vp16,14UAS:Clover)UoBL1* line. One stable transgenic line per docking line was generated with the miR-430 reporter construct. Expression driven by the miR-430 promoter was detected in both lines at 256/512-cell and sphere stages by RT-qPCR (**Figure 3B**). This expression, together with the lack of detectable expression of the transgene at earlier or later stages, is in line with the expression dynamics of the endogenous miR-430 cluster. In contrast, reporter expression from the γ-crystalline promoter known to be active at later stages in the developing lens (Davidson *et al.*, 2003) was detected only at day 4 (96 hpf) as expected, indicating the transcriptional competence of the integration site. These results together demonstrate that the mirR-430 proximal promoter sequence contains sufficient regulatory information to activate transcription at early pre-MBT/ MBT stages.

### The miR-430 gene cluster and its activity are required for the formation of a transcription body at the minor wave of genome activation

To dissect the contribution of underlying genomic sequence and transcription from the miR-430 chromosome 4 cluster to the formation of the transcription body, we used the CRISPR system to specifically target the miR-430 loci and recruit Cas9 protein-associated loss of function tools onto the genome at the core of the transcription body. This system was optimised for both lesion generating active Cas9, which can induce double strand DNA breaks (DSB) in the targeted genomic loci, and with catalytically dead Cas9 (dCas9), which can bind target sequences, without generating lesions and blocks transcriptional by sterically hindering binding and progression of Pol II, in a process termed CRISPRi (Larson *et al.*, 2013).

To address the impact of Cas9-mediated lesion and dCas9 inhibition on miR-430 activity, we used quantitative PCR analysis of miR-430 expression. A dramatic reduction in miR-430 RNA was detected at the 512-cell stage, following targeting of the miR-430 cluster by either active Cas9 or dCas9, comparable to total transcription loss achieved through global transcription inhibitor treatment (triptolide or α-amanitin) (**Figure 4A**).

**Figure 4.**
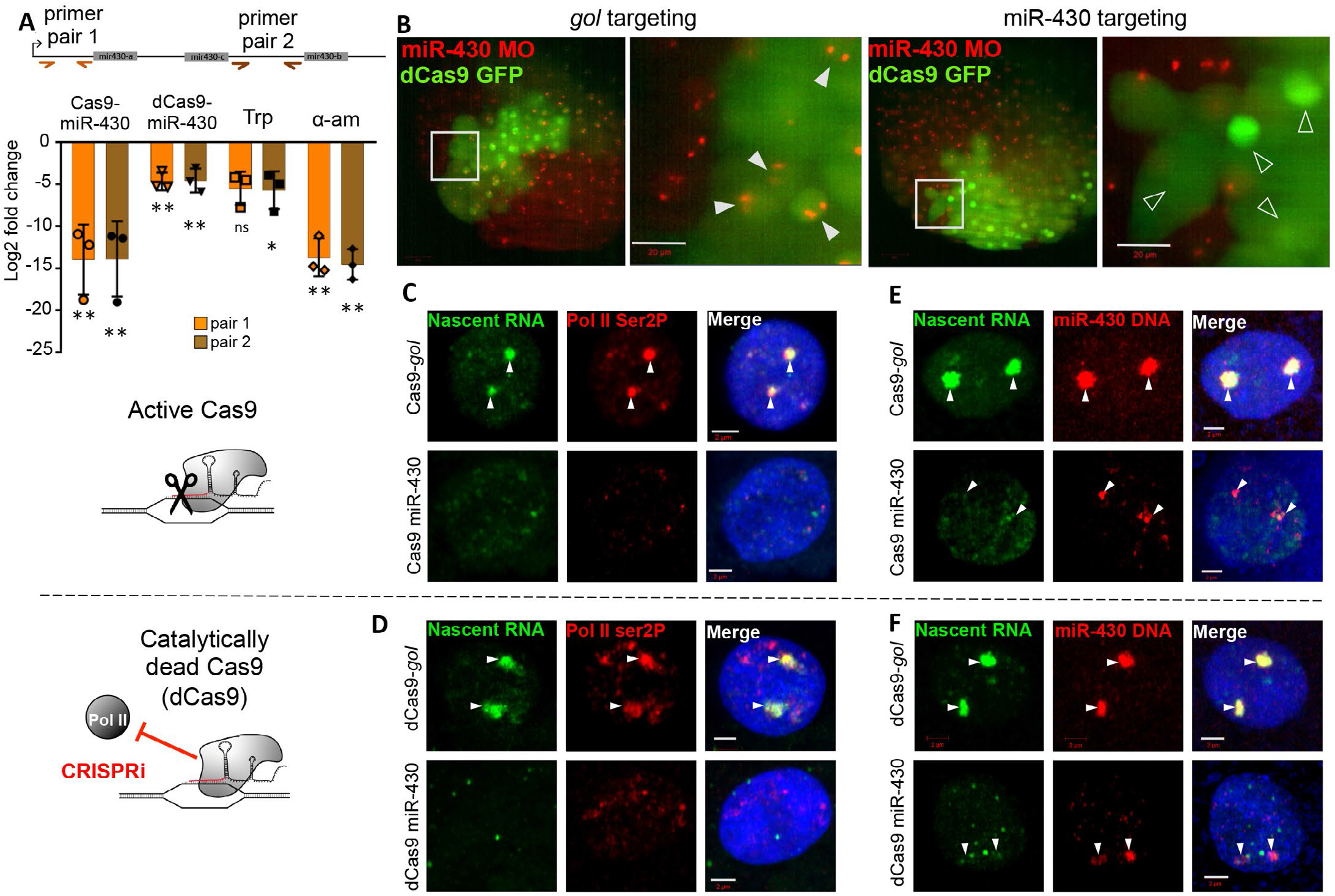
Manipulations that limit transcription of the miR-430 cluster also disrupt the transcription organiser. **A,** Schematic of a single miR-430 triplet repeat showing the position of two RT-qPCR primer sets (orange and brown arrows). Chart shows relative levels of miR-430 RNA at the 512-cell stage, normalised to 5S rRNA, following manipulation by miR-430 promoter targeting by active lesion generating Cas9 or catalytically dead-Cas9 (dCas9), and triptolide (Trp) or α-amanitin (α-am) treatment, compared to control manipulations of active Cas9 targeting *gol* splice junction, dCas9 targeting *gol* splice junction and injection control, respectively. Data is based on three biological repeats. **B,** Live cell images of miR-430 MO (red) labelled transcription bodies in 512-cell stage embryos following either miR-430 promoter or *gol* splice junction mosaic targeting with dCas9-GFP (green). Grey boxes indicate enlarged region shown on the right; *gol* targeted: 10 embryos, miR-430 targeted: 8 embryos. **C,D,** EU labelling of nascent RNA(green),Pol II ser2P(red) immunostaining and DAPI (blue) in 512-cell stage embryos. White arrows indicate remnants of the transcription body. (C)Active Cas9; *gol* targeted: 6 embryos, 31 nuclei. miR-430 targeted: 6 embryos, 28 nuclei. (D)Catalytically dead Cas9; *gol* targeted: 8 embryos, 52 nuclei. miR-430 targeted: 5 embryos, 29 nuclei. **E,F,** EU labelling of nascent RNA (green) combined with FISH for miR-430 DNA (red) and DAPI (blue) at the 512-cell stage. (E) Active Cas9; *gol* targeted: 4 embryos, 21 nuclei. miR-430 targeted: 4 embryos, 41 nuclei. (F) Catalytically dead Cas9; *gol* targeted: 5 embryos, 41 nuclei. miR-430 targeted: 8 embryos, 66 nuclei.

Using the MoVIE technique of *in vivo* imaging of miR-430 nascent RNA accumulation in living embryos (Hadzhiev *et al.*, 2019), we monitored miR-430 expression together with *in vivo* detection of green fluorescent protein fused to dCas9, to determine the effect of miR-430 CRISPRi on the accumulation of miR-430 RNA. Single cell stage injection of miR-430 targeting labelled morpholinos, was combined with a second injection into an individual cell at the 8-cell stage, delivering dCas9-GFP and guide RNAs targeting either miR-430 or a control gene (*gol*) in a mosaic fashion. This dual injection setup allowed for the ubiquitous labelling of miR-430 RNA across the embryo from the 64-cell stage, alongside mosaic and trackable delivery of CRISPRi reagents (**Figure 4B**). Using these imaging tools, we observed that loss of miR-430 transcription was throughout the cell cycle and specifically occurred in cells containing dCas9 knock-down reagents targeting the miR-430 cluster, suggesting a cell autonomous direct effect (**Figure 4B, Supplementary Figure 4A, B, Supplementary movie 1 & 2**). The complete loss of miR-430 morpholino labelled foci in GFP^+^ cells specifically following miR-430 targeting, shows that this CRISPRi is effective despite the highly repetitive nature of the miR-430 cluster **(Figure 4B).**

To establish the role of the miR-430 locus in the formation of the transcription body, through the enrichment of elongating Pol II (labelled with an anti-Pol II CTD Ser2P antibody) at the chromosome 4 miR-430 cluster, we used Cas9 lesions to disrupt the miR-430 locus. In 512-cell stage embryos where the Cas9 disruption was targeted to the miR-430 locus (**Figure 4C,E**), but not to a control locus (*gol*), we observed a loss of the transcription body; with nascent RNA accumulation and Pol II Ser2P signal reduced to background levels, which was accompanied by a reduction in the size of miR-430 territory detected by DNA-FISH (Figure 4E). To exclude that the loss of the transcription body is due to potential chromosomal rearrangements caused by multiple Cas9 lesions, or complete loss of the miR-430 cluster, we repeated the transcription body disruption analysis with dCas9 targeting to the miR-430 cluster, which inhibits transcription of the miR-430 locus, but lacks legion-generating catalytic activity. Again, we observed a loss of the transcription body and a reduction in the size of miR-430 territory (**Figure 4D,F**). Global transcription inhibition by either triptolide or α-amanitin, also resulted in a loss of the transcription body and reduction in miR-430 DNA-FISH signal volume (**Supplementary Figure 4C-H**).

Together these observations suggest that transcription of the miR-430 cluster is required for the formation and maintenance of the transcription body and suggests a transcription organiser role for miR-430. We show that the expansion of the transcribed territory in 3D space is dependent on miR-430 transcription, in line with previous models where RNA accumulates around the repeats as they are transcribed forming a growing transcription bubble, which expands the volume occupied by the loci (Hilbert *et al.*, 2021). Our experiments identify high promoter density and subsequent transcription of the 0.6 Mb locus of miR-430 repeats as the primary reason for this transcription body formation.

### The miR-430 locus acts as a transcriptional organiser and influences early zygotic transcription of znf genes on the long arm of chromosome 4

Our nascent RNA and Pol II Ser2P imaging of pre-ZGA embryos shows that most of the minor wave transcription and nascent RNA accumulation is concentrated within the transcription body, associated with miR-430. This led us to query whether other early transcribed loci contributed or in some way interacted with the transcription body, to enable their early escape from global transcription repression. To test whether the transcription body had a role in nuclear organisation during the minor wave of ZGA, we investigated whether manipulation of miR-430 or its transcriptional activity would impact on the first wave activity of early expressed genes on chromosome 4 and on other chromosomes. To this end first we identified the location and activity dynamics of minor wave active genes on chr 4 and on other chromosomes.

The miR-430 cluster sits in a unique genomic environment, relatively close (1.7 Mb) to the centromere of chromosome 4, flanking a 40 Mb gene-poor region that stretches for most of the long arm of chromosome 4 and shows distinct transcriptional dynamics from the rest of the genome (White *et al.*, 2017) (**Figure 5A**). We and others have previously shown that the long arm of chromosome 4, despite being relatively gene poor contains a large number of *znf* genes mainly encoding C2H2 family proteins, some of which show early zygotic transcription (White *et al.*, 2017). We also investigated the transcriptional dynamics of the *znf* genes contained within the long arm, of chromosome 4, neighbouring the miR-430 cluster, which we have previously shown to colocalise their RNA with that of miR-430 genes (Hadzhiev *et al.*, 2019). Our CAGE-seq (Nepal *et al.*, 2013) and nascent RNA-seq (Heyn *et al.*, 2014) analyses revealed different transcriptional behaviours relating to distinct promoter architectures within these *znf* genes. A subset of these *znf* genes is expressed at the minor wave of ZGA (**Figure 5A-C**). and have a sharp promoter type with a TATA-box. This sharp promoter structure of the early expressed chromosome 4 *znfs* is shared with other minor wave genes (**Figure 2A**) and suggest that distinct transcriptional machinery may recognise these sharp promoters to drive early zygotic transcription. In contrast, additional *znf* genes intermingled with the sharp promoter *znf*s along the long arm of chromosome 4, are not expressed at the minor, but at the major wave of ZGA and have broad promoter type (**Figure 5A-C**).

**Figure 5.**
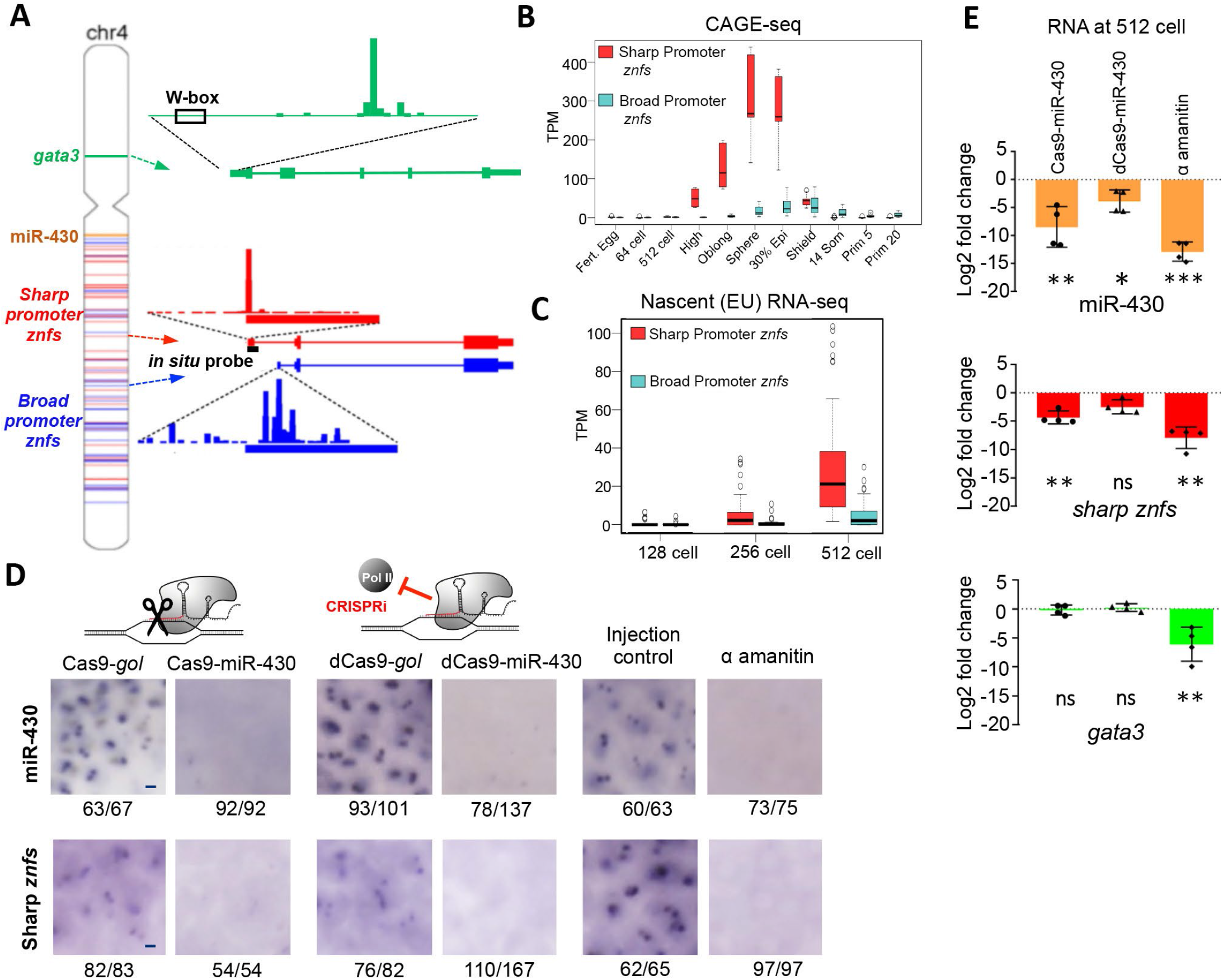
MiR-430 facilitates early zygotic transcription of minor wave znf geneson the chromosome 4 long arm but not of other minor wave genes on chromosome 4 short arm or on other chromosomes. **A,** Location of gene of interest on chromosome 4: *gata3*, miR-430 cluster plus broad and sharp promoter *znfs* as blue and red bands, respectively. Right: Promoter architecture from sphere stage CAGE-seq. **B,** Promoter activity during early development by CAGE-seq for chromosome 4 sharp and other promoter znfs. **C,** Gene expression levels from EU-seq for sharp and other promoter znfs from chromosome 4, during ZGA. **D,** Nuclear pairs of nascent RNA foci (arrowheads) detected by chromogenic ISH of miR-430 and sharp promoter *znf* genes at 512-cell following manipulations. Numbers indicate frequency of staining pattern, scale bar = 5μm. **E,** Gene expression analysis, by RT-qPCR of 512-cell stage embryos, shown as mean log2 fold change of 4 biological repeats. Error bars represent standard deviation.

To determine the extent of influence of the transcription body on minor wave of ZGA we investigated the response of early transcribed *znf* genes and two single copy minor wave genes selected (Heyn *et al.*, 2014)): *klf17* on chromosome 2 and *gata3* on the short arm of chromosome 4 to loss or block of the miR-430 locus. The minor wave activation of *klf17* and *gata3* was confirmed by CAGE-seq showing detectable signal as early as 512-cell stage, one cell cycle before ZGA (**Supplementary Figure 5A**). We observed a robust reduction in miR-430 RNA levels in response to miR-430 targeting by Cas9 or global transcription block by α-amanitin, and a modest reduction in miR-430 RNA in response to dCas9 targeting the miR-430 cluster (**Figure 5D,E**, **Supplementary Figure 5C**). Following miR-430 targeted manipulations and global transcription block we observed consistent reduction in RNA from chromosome 4 minor wave sharp promoter *znf*s by RT-qPCR, which was also confirmed by *in situ* hybridisation in whole embryos (**Figure 5D,E**, **Supplementary Figure 5C**). Degree of miR-430 signal reduction correlated with loss of early sharp *znf* expression at the 512-cell stage. This suggests that the transcription body facilitates early zygotic transcription initiation from sharp promoter *znf*s with shared promoter architecture properties similar to miR-430 and contained within the long arm of chr4, and in good agreement with the previous observation of colocalization of their RNAs in a transcription body (Hadzhiev *et al.*, 2019).

Next, we asked whether miR-430 activity and the transcription body influence early transcribed genes on the same chromosome but separated by the centromere. We identified low but detectable *gata3* transcription from the 512-stage in our CAGE-seq analysis (**Supplementary Figure 5A**) confirming previous reports of its early activity (Heyn *et al.*, 2014). This single copy gene is located on the short arm of chromosome 4, has a sharp promoter structure (**Figure 5A**) and is one of the nearest annotated genes to the centromere, making it an ideal target to investigate the spread of influence from the transcription body associated with miR-430. At the 512-cell stage, following miR-430 targeting by either Cas9 or dCas9 protein, *gata3* RNA levels showed no significant difference from *gol* targeted controls (**Figure 5E**), while global transcription inhibition greatly reduces *gata3 RNA* confirming its zygotic expression at this stage in the absence of manipulation. The lack of *gata3* early transcriptional response suggests that the mechanisms facilitating early zygotic transcription from sharp promoter types are not restricted to the long arm of chr 4, and that mir430 activity is not required for all genes active on chr 4 during the minor wave of ZGA, probably because they are not interacting with the transcription body.

We then asked whether loci or transcripts from other chromosomes also interacted with miR-430 the transcription body, focussing on *klf17*, a single copy gene residing on chr2, which shows transcriptional activity at the 512-cell stage and a sharp promoter structure with TATA-box (**Figure 6A**) To address co-localisation of loci and RNA we used nascent RNA labelling combined with DNA or RNA FISH to identify the sub-nuclear topological relationship between the transcription body at the miR-430 locus and *klf17.* We observed that the pair of *klf17* loci reside in a distinct chromosomal territory to the transcription body, marked by either nascent RNA or by the miR-430 loci itself, at the 512-cell stage (92.4% and 95.1% of *klf17* loci are topologically distal to a transcription body, respectively) (**Figure 6B,C**). Interestingly, nascent RNA can sometimes be observed accumulating proximal to the *klf17* loci (white arrows **Figure 5H**), confirming *klf17*’s zygotic transcription at this stage, plus also emphasising the small number of transcriptionally active loci across the whole genome and their relative expression level to the miR-430 loci, at this early stage. This data shows that although the transcription body is highly enriched for nascent RNA it is not a unique site of transcription within the early nucleus and that single copy genes can escape global transcriptional repression despite occupying distinct topological domains from the transcription body.

**Figure 6.**
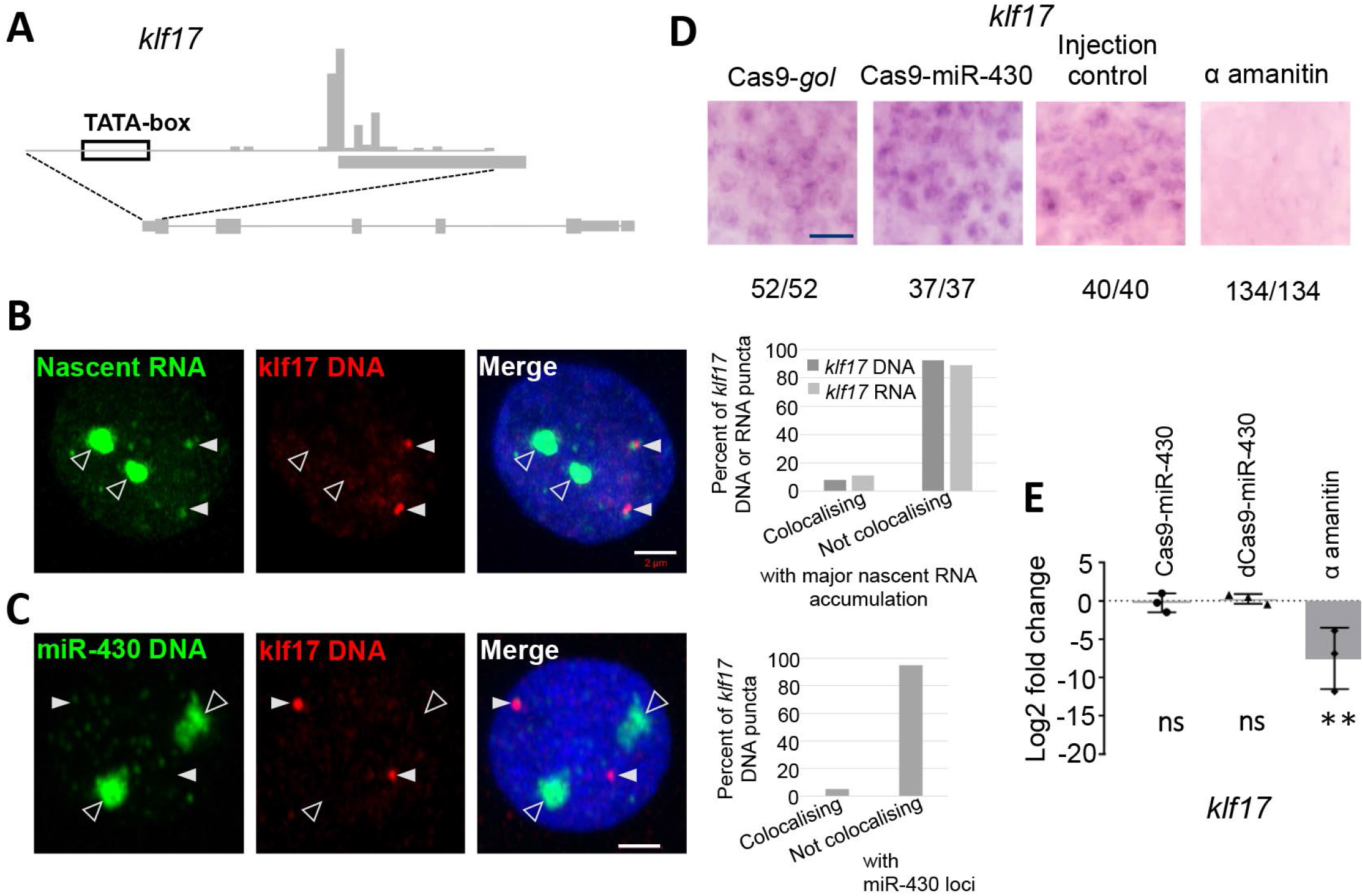
MiR-430 does not influence early zygotic transcription of minor wave genes on other chromosomes. **A,** Promoter architecture from sphere stage CAGE-seq for *klf17*. **B,** Nascent RNA (green) combined with FISH for *klf17* DNA (red) and DAPI (blue) at 512-cell. 59 nuclei from 16 embryos. Right, chart shows frequency of colocalization between klf17 DNA or RNA and major nascent RNA accumulation (transcription body) in 3D space, (59 and 54 nuclei respectively).**C,** Double DNA-FISH labelling the miR-430 cluster (green) and the *klf17* locus on chromosome 2 (red) at 512-cell stage counter-stained with DAPI (blue). 51 nuclei from 8 embryos. Chart shows frequency of colocalization between *klf17* DNA and miR-430 DNA in 3D space. **D,** Nascent *klf17* RNA staining at 512-cell following manipulations. Numbers indicate frequency of staining pattern, scale bars = 25μm. **E,** *klf17* RNA level analysis, by RT-qPCR of PCNA synchronised 512-cell stage embryos following various manipulations. Results shown are mean log2 fold change of 3 biological repeats. Error bars represent standard deviation.

These fixed imaging approaches highlight the dynamic nature of early genome topology and could not exclude the possibility of a highly transient interaction between early expressed loci around the genome and the transcription body at the chromosome 4 miR-430 cluster. To investigate whether any transient interaction enabled miR-430 activity to influence early *klf17* transcription we followed *klf17* RNA levels upon miR-430 locus and locus activity disruption. The manipulation of miR-430 by either Cas9 or dCas9 did not have significant effect on *klf17* RNA levels, while global transcription inhibition induced a significant loss in *klf17* RNA (**Figure 6D,E**). Other early zygotically expressed genes, *wnt11f2* and *hspb1* (both chr5), also failed to show a response to miR-430 targeted manipulations (data not shown). This lack of response to miR-430 manipulations suggests that the transcription body does not exert a global effect influencing minor wave activation of genes on other chromosomes, or on the short arm of chromosome 4 (*gata3*).

## DISCUSSION

Much is already known about the mechanisms and regulatory networks underlying the main wave of zygotic genome activation, but the mechanisms governing the very first steps of global genome activation are still unclear (Lee *et al.*, 2014; Schulz and Harrison, 2019; Vastenhouw *et al.*, 2019). Here we explored the genetic mechanisms underlying the activation of minor wave genes in zebrafish. We chose to study the miR-430 gene cluster, which are the first known expressed genes in zebrafish (Heyn *et al.*, 2014). Our results show that this gene cluster is composed of unexpectedly high promoter/gene density, which define and is required for the formation of an unique transcription body. This transcription body is enriched for nascent RNA and Pol II Ser2P during pre-ZGA cell cycles (Chan *et al.*, 2019; Hadzhiev *et al.*, 2019; Hilbert *et al.*, 2021) including not only miR430 gene products but that of a set of *znf* genes residing on the same chromosome 4 long arm where miR-430 cluster is located. Thus, we propose that the miR-430 cluster functions as a transcriptional organiser for a set of minor wave genes occupying the same chromosomal territory. Moreover, we have demonstrated that minor wave genes including the miR-430 cluster share epigenetic and promoter architecture features, which seem to contribute to their selective activation before that of the rest of the genome (Figure 6).

### Extreme density of promoter/gene cluster underlies the formation of a minor wave-associated transcription body

We have previously demonstrated that miR-430 nascent RNA (primary microRNA) products accumulate in a unique transcription body during the first wave of genome activation in zebrafish. We demonstrated by several independent lines of evidence that the copy number of miR-430 transcribed units is almost two orders of magnitude higher than described in the presently available reference genome GRCz11. This observation highlights the importance of generation of end-to-end genome assemblies, which are necessary to resolve highly repeated loci, a challenging task by conventional genome sequencing tools (Treangen and Salzberg, 2011). A long-read zebrafish genome sequence was also reported recently (Yang *et al.*, 2020). Our manual analysis of that published chromosome 4 assembly calculated ~40 promoters. In our assemblies we show substantially larger locus, which we achieved by several independent analyses and more than one zebrafish strains suggesting that the hundreds of copies of promoters we detect is a robust feature of common zebrafish strains. Interestingly, miR-430 homolog loci are also multicopy in several teleosts like medaka and goldfish (manual analysis of improved assemblies of the medaka genome (Ichikawa *et al.*, 2017) and *de novo* assembled goldfish genome (Chen *et al.*, 2019)), likely indicating a crucial role for these miRNAs in clearance of maternal mRNAs (Giraldez *et al.*, 2006). Nevertheless, the exact composition of the gene sets within the cluster appears to be variable and merits further analysis given the genetic significance of zebrafish chromosome 4, with unique gene organisation (White *et al.*, 2017) and implication in sex determination (Yamazaki *et al.*, 1989)

Importantly, genome activation is also concentrated in discrete, Pol II Ser2P-containing nuclear compartments in *Drosophila* (Blythe and Wieschaus, 2015; Chen *et al.*, 2013; Hug *et al.*, 2017). Similarly to zebrafish, genome activation in human and mouse embryos involves multicopy transcribed repeats (Grow *et al.*, 2015; Macfarlan *et al.*, 2012). Recent *in vivo* work with transcription imaging technologies has revealed distinct Pol II Ser2P accumulation foci in several mammalian cells (Cisse *et al.*, 2013; Conic *et al.*, 2018) indicating formation of sizeable transcription hubs, which are reminiscent to that seen in an exaggerated form in the zebrafish embryo. Overall, these observations from a variety of models including mammals suggest that gene clusters are often organised into distinct nuclear topologies (Cook and Marenduzzo, 2018). Thus, understanding the mechanisms underlying the formation of the zebrafish transcription body may reveal genome activation-associated transcription control as well as fundamental regulatory principles of transcription-associated nuclear topology formation relevant also in mammals.

The 0.6 Mb miR-430 region packed with about 300 promoters suggested that the mechanisms for the early escape from global transcriptional repression may arise from bulk properties and macro nuclear dynamics. The miR-430 promoter-dense cluster may form a transcriptionally competent bubble within which the nucleocytoplasmic ratio can be passed in the local microenvironment. However, this model is not supported by two observations. Firstly, miR-430 early activation does not respond to change of the nucleocytoplasmic ratio by ploidy manipulation (Chan *et al.*, 2019; Hadzhiev *et al.*, 2019). Secondly, in this study, minor wave activity of a single miR-430 promoter inserted in a heterologous genomic location was demonstrated. This indicates that the large copy number of miR-430 was not necessary for reaching a threshold of transcriptionally competent state, but that the promoter sequence contained the necessary regulatory information for early activation. We propose that the very high promoter density enables the formation of a transcriptionally favourable microenvironment around the miR-430 locus, which facilitates the formation of nuclear body, which may further boost the promoter-autonomous capacity for responsiveness of the miR-430 promoter and other gene promoters occupying the transcription body (Figure 7).

**Figure 7.**
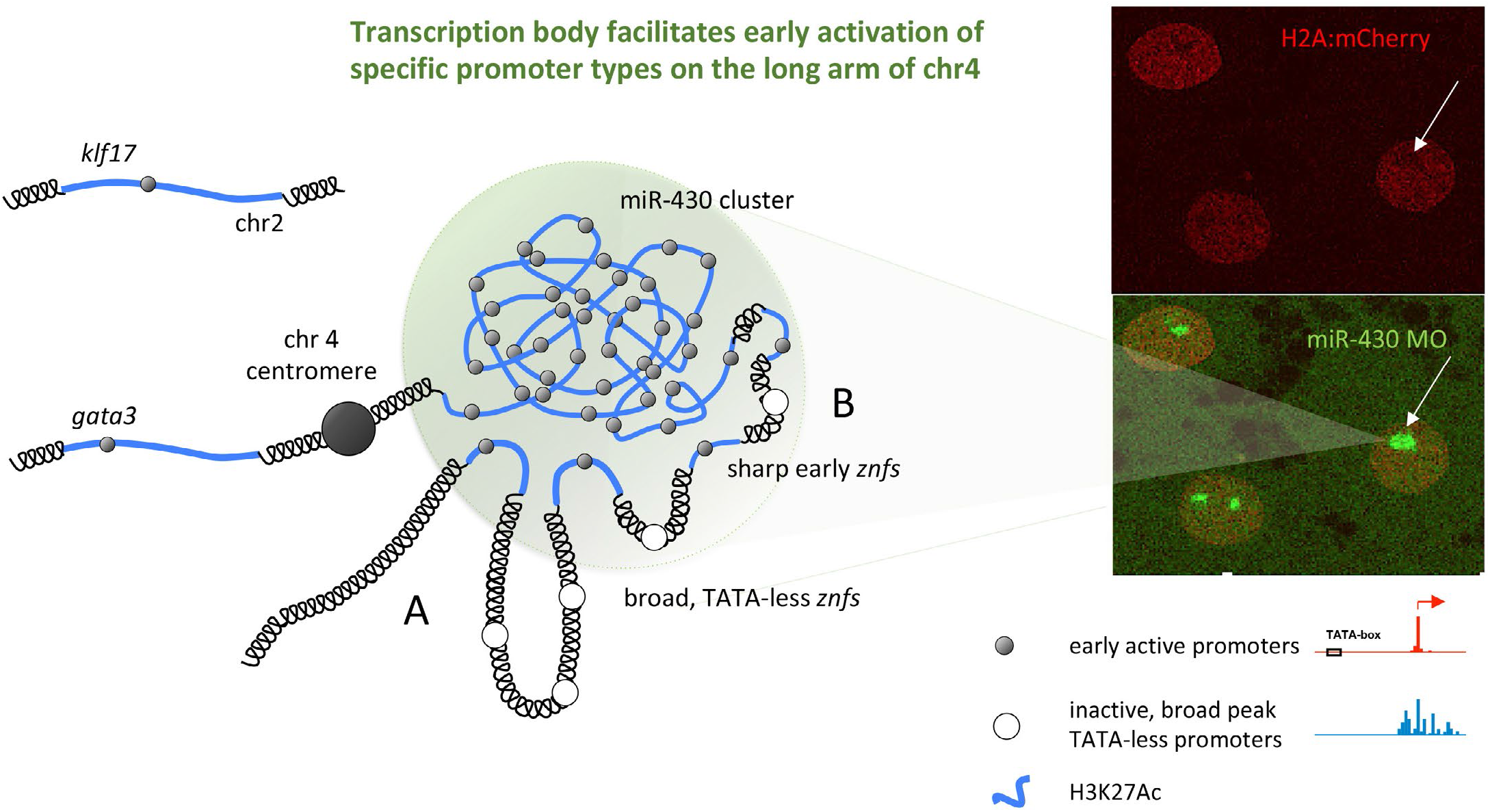
Model for miR-430 cluster acting as transcription organiser in forming a transcription body, which facilitates transcription of genes on the long arm of chromosome 4. Two models of potential action of the transcription body are shown. **A** The miR-430 cluster with extreme promoter density forms amicroenvironment, which facilitates minor wave activity of miR-430 genes and additional genes of the chr 4 long arm. The *znf* genes, which share promoter architecture features with miR-430 genes are colocalising and are activated in the transcription hub. *Znf* genes with distinct promoter features are expelled from the transcriptionally active hub formation. **B**, The miR-430 cluster facilitates activation of promoters with the transcription body, which share promoter architecture features, while colocalising promoters that have distinct promoter features are not recognised or not activated by the transcription machinery active in the transcription body due to sequence incompatibility. In both models minor wave gene activation can also occur distal to the transcription body, such as at the *gata3* and *klf17* loci

### Distinct promoter features shared by minor wave ZGA genes

Previous studies have shown that miR-430 transcription requires H3K27ac deposition, binding of stem cell factors Sox19b/ SoxB1 Nanog and Pou5f1/Pou5f3 (Miao *et al.*, 2020), and it is characterised by CTCF and Rad21 deployment (Chan *et al.*, 2019; Meier *et al.*, 2018; Miao *et al.*, 2020) at a stage when topology associated domains have not yet formed (Kaaij *et al.*, 2018; Wike *et al.*, 2021). However, the question to answer remains: what is the mechanism which triggers miR-430 transcription and that of other minor wave transcribed genes? While the stem cell factors Pou5f3/Pou5f1, SoxB1/Sox19b and Nanog were previously shown to be required for miR-430 gene activation (Lee *et al.*, 2013; Miao *et al.*, 2020), they are unlikely to selectively regulate minor wave genome activation, as a large number of major wave genes also depend on these factors (Gao *et al.*, 2020; Lee *et al.*, 2013; Miao *et al.*, 2020; Palfy *et al.*, 2020; Veil *et al.*, 2019).

When comparing promoter features of minor wave genes to other zygotic-active genes we observed a lack of H3K4me3 at miR-430 and minor wave genes. This histone mark is generally associated with gene regulation and has been shown to be present on promoters preceding genome activation and even detected in sperm (Haberle *et al.*, 2014; Lindeman *et al.*, 2011; Wu *et al.*, 2011; Zhu *et al.*, 2019) While this histone mark may be paternally inherited as seen in mouse (Lismer *et al.*, 2021), our findings argue against these mechanisms being involved in activation of miR-430 and other minor ZGA genes in zebrafish. However, we demonstrated that that minor wave genes are enriched in sharp promoters, carry TATA-box (similarly to *Drosophila* (Chen *et al.*, 2013)), both features often associated with lack of H3K4me3 (Reviewed in (Sandelin *et al.*, 2007)). TATA-dependent sharp promoters are distinctly lacking in promoters of the major wave genes (Haberle *et al.*, 2014) suggesting important distinction and likely distinct transcription initiation machinery acting in them (Haberle *et al.*, 2019; Levine *et al.*, 2014; Muller *et al.*, 2010). However, TATA-dependent sharp promoters are not restricted to the minor wave and yet unidentified factors may also play crucial roles in specifying early zygotic gene activity. Future analysis of minor wave regulatory elements and their occupancy would aid the identification of as yet unidentified motifs and chromatin opening or pioneering factors (recent examples for such factors include GAF in Drosophila (Gaskill *et al.*, 2021) or Dux in mammals (De Iaco *et al.*, 2017; De Iaco *et al.*, 2020; Hendrickson *et al.*, 2017), which play key roles in these early mechanisms.

### MiR-430 promoter cluster as transcription organiser in the transcription body

Previously it was demonstrated that miR-430 expression is a component of the transcription body (Chan *et al.*, 2019; Hadzhiev *et al.*, 2019; Hilbert *et al.*, 2021). Here we demonstrate that transcription of the miR-430 cluster is specifically required for the formation of this transcription body. Similarly, we observe a reduction in nuclear volume occupied by the miR-430 cluster of chromosome 4 following CRISPRi targeting. This correlates with the proposal that high levels of transcription expand chromatin (Hilbert *et al.*, 2021), and form a favourable microenvironment, which further promotes miR-430 transcription. An additional role of the transcription body may be to provide favourable chromatin environment for efficient facilitation of transcription-associated processes, such as co-transcriptional Drosha-dependent primary miRNA precursor processing of miRNAs (Ballarino *et al.*, 2009). Future experiments could distinguish contributions of nascent RNA production from RNA processing in the process of transcription chromatin decondensation/phase separation in the transcription body.

We demonstrated through the analysis of minor wave ZGA genes that the highly dynamic nuclear body formed by miR-430 activity influences the early transcriptional activity of *znf* genes, contained within the long arm of chromosome 4. We showed that these *znf* genes selectively share a specific promoter structure including lack of H3K4me3, presence of the TATA box and sharp transcription initiation profile (**Figure 6**). Importantly, the effect of miR-430 is limited to the chr 4 long arm and other minor wave genes appear to be activated independently on the short arm of chr 4 or on other chromosomes. These observations lead to a model, whereby high-density promoter cluster of miR-430 facilitates the formation of a transcription body encompassing chr 4 activity domains. The high density of binding sites for stem cell factors and yet unidentified additional transcription factors, contained within the miR-430 promoter cluster, function as ‘enhancers’ for genes, with enhancer interaction-compatible promoter architectures on chr 4 long arm (**Figure 7**). In this context the 0.6 Mb miR-430 locus enriched in H3K27ac and stem cell factor binding sites is not dissimilar from what previously was described as an enhancer cluster or also called super-enhancer (Hnisz *et al.*, 2017). The enhancer effect of the large number of promoters observed here fits also with a recent computational model of enhancer and promoter interactions, which suggest generality of equivalent topological and functional impact by enhancer or promoter clusters (Zhu *et al.*, 2021).

Importantly, a large number of *znf* genes interspersed with minor wave active *znf*s do not get activated before the main wave of ZGA despite being scattered along the chr 4 long arm and differ in their promoter architecture (broad promoter, lack of TATA box). These observations suggest that the enhancing or transcription organising effect of miR-430 locus is limited to genes with compatible promoter architectures. This observation either indicates action of promoter-enhancer interaction specificity mechanisms defined by core promoter sequence determinants (Gehrig *et al.*, 2009; Haberle and Stark, 2018; Zabidi *et al.*, 2015) or that major wave ZGA genes with distinct promoter architectures do not respond to specific pioneering transcription factors. In the later case, these pioneering factors may locally function at minor wave gene promoters and likely interact with general transcription factors selectively binding the TATA-box, but fail to act upon different promoter architectures present in proximity. An alternative mechanism for the local influence of the transcription body could be spreading of an epigenetic signal along the long arm of chromosome 4, permitting increased chromatin accessibility. This increased chromatin accessability may be regulated through the activation of transposable elements, which have been shown, in turn, to regulate zinc finger proteins similar to those enriched on chr 4 and active during the minor wave in zebrafish (Turelli *et al.*, 2014).

Future studies will need to address whether this transcription body acts by enriching for key general and specific transcription factors, which may selectively recognise TATA box-dependent, sharp TSS core promoter codes or provide favourable chromatin environment exploited by genes on chr 4 sharing a chromosome territory with miR-430 gene cluster

## METHODS

### Zebrafish maintenance

All zebrafish strains were maintained in a designated facility (according to UK Home Office regulations) in a recirculating system (ZebTEC, Tecniplast) at 26°C in a 10-hour dark, 14-hour light photoperiod and fed 3 times daily.

Zebrafish experiments were restricted to embryos in early developmental stages and adults were only used for natural breeding. Animal work presented in this study was carried out under the project licence P51AB7F76 assigned to the University of Birmingham, UK

Zebrafish embryos were obtained by putting together a pair of zebrafish (male and female separated by a divider) in a specifically designed crossing cage in the evening. On the next morning after the start of the light period, the dividers were removed, bringing the two sexes together which initiates spawning. Embryos were collected shortly (5-15 min) after laying by using a small net or tea strainer.

For the PacBio assembly, Gynogenic offspring were generated as described in The zebrafish book (Westerfield and ZFIN., 2000). Homozygosity was confirmed by PCR amplifying 8 independent loci and confirming there were no SNV’s within any sequence.

### Isolation of HMW zebrafish genomic DNA

High molecular weight genomic DNA was isolated from one AB male by proteinase K and phenol extraction (Green and Sambrook, 2017). Adult fish were euthanized by overdose of anaesthetic (Tricaine) and snap frozen in liquid nitrogen. The frozen tissue was pulverised with liquid nitrogen precooled ceramic pestle and mortar. The powder was slowly spread over the surface of 20 ml lysis buffer in 100 ml beaker. After complete submerging of the tissue powder into the lysis buffer the lysate was transferred to 50 ml falcon tube and treated with pancreatic RNase for 1 hour at 37 °C and proteinase K overnight at 50 °C, followed by three phenol extractions. The DNA was precipitated from the purified lysate by addition of 0.2 volume of 10 M ammonium acetate and 2 volumes of ethanol, washed twice with 70% ethanol air-dried and reconstituted in TE buffer.

### Oxford Nanopore sequencing

Nanopore sequencing library was prepared with Genomic DNA Ligation Sequencing Kit (Oxford Nanopore, SQK-LSK110) according to the manufacture instructions, using 1 μg genomic DNA as input and sequenced on Oxford Nanopore Promethion device. The sequencing run resulted in 1.7133×10^7^ reads and 1.3273×10^11^ bp in total (app. 44x coverage of the ~1.7Gb zebrafish genome).

### Pacific Biosciences (PacBio) sequencing

Genomic DNA was purified using TissueLyser II (Qiagen) and Blood & Cell Culture DNA Maxi Kit (Qiagen). The molecular size of genomic DNA at the peak of 40- to 50-kb was confirmed using the Pippin pulse electroporation system (NIPPON genetics). Genomic DNA from the TU/AB double heterozygous fish described above was used to perform whole-genome shotgun sequencing on a PacBio RS II sequencer. ≈16 million subreads with a peak length of ≈8kb (100X covergage).

### *De novo* genome assembly of long read data

The Oxford nanopore and PacBio sequencing reads were assembled into contigs using Canu software v.2.1.1 (Koren *et al.*, 2017) with the following parameters for nanopore reads: `-nanopore genomeSize = 1.7g minReadLength = 5000 minOverlapLength = 1000` and PacPio reads: `-pacbio genomeSize =1.7g minReadLength =1000 minOverlapLength = 500`

To produce guided whole genome assembly from the nanopore data a haplotype purge was performed on the assembled contigs with purge_dups (Guan *et al.*, 2020), following the pipeline guide with default parameters as outlined on (https://github.com/dfguan/purge_dups)

A guided (using the current zebrafish genome (GRCz11) as reference) scaffolding into chromosomes of the haplotig purged assembly (~1.4 Mb in size) was performed with RaGOO v1.1 (Alonge *et al.*, 2019) according to the default pipeline (https://github.com/malonge/RaGOO) enabling misassembly correction with the error corrected nanopore reads (generated by Canu during the contig assembly step):`ragoo.py hap_purged_ctigs.fa GRCz11chr_ref.fa -R CorrNPRds.fa -T corr`.All assemblies were carried out on the University of Birmingham high performance computing cluster BlueBEAR.

### Estimation of miR-430 promoter number form raw long reads

To estimate the miR-430 promoter copy number a 250 bp promoter region sequences of miR-430 and 6 single copy genes. The single copy genes were randomly selected from the Benchmarking Universal Single-Copy Orthologs (BUSCO) database (Actinopterygii gene set) and confirmed by BLAST search that the selected 250 bp sequence produced a single hit in the current (GRCz11) zebrafish reference genome.

The miR-430 and the 6 single copy genes promoter sequences were used to perform a BLAST search against raw reads BLAST database for each sequencing run. BLAST hits with bit score >= 150 were counted for each gene and miR-430 counts were normalised to each of the 6 single copy genes and the average was used as miR-430 promoter number estimate.

### Estimation of miR-430 promoter number by qPCR

To estimate the miR-430 promoter copy number by qPCR using the standard curve absolute quantification method genomic DNA from three stains AB, Tuebingen (TU) and Tuebingen Longfin (TL) was prepared from 50, 2 day old embryos collected from a single pair for each strain. The DNA extraction was carried out with the PureLink™ Genomic DNA Mini Kit (ThermoFisher Scientific, K182001) following the manufacture instructions.

A plasmid containing a single copy of the miR-430 promoter was used to generate dilution series (8 in total) simulating presence of 2 to 256 copies of the promoter sequence. The dilution series were used to generate standard curves and were supplemented with 50 ng human genomic DNA (which does not contain the target sequence) to mimic genomic template complexity of the samples in the PCR reaction. 50 ng of genomic DNA from each strain was used as template in a PCR reaction with 0.5 μM of each primer (FP: AGGGTGTGAGTTTGCATGAGAGTG; RP: GGCTTTATGTGTCCTGCAGCAG) and PowerUp SYBR Green Master Mix (ThermoFisher Scientific, A25742). All reactions were ran in triplicates.

### Promoter density analysis

CAGE-seq data from Nepal et. al (Nepal *et al.*, 2013) were mapped to the zebrafish long-reads assembly genome with Bowtie v1.2.3 (Langmead *et al.*, 2009) in default n-mode allowing 2 mismatches in the 27 bp seed region and reporting all the best alignments up to 1000 (`-S -n 2 -l 27 -a -m 1000 --best – strata`). The CAGEr package (Haberle *et al.*, 2015) was used for downstream processing of the data. First, CAGE Transcription Start Site (CTSS) were called in each of the analysed samples removing the additional G nucleotide (due to the CAGE protocol) not mapping to the genome. Next, reads were counted at each CTSS and subsequently normalized as tags per million (tpm). CTSS supported by at least 0.5 tpm in at least two samples and closer than 20 nucleotides were clustered thus defining the so-called transcriptional clusters (TCs), discarding singletons TCs (i.e., clusters containing only one CTSS). TCs were then trimmed on the edges to obtain a more robust estimation of the promoter width. Toward this end, first the cumulative distribution of CAGE signal along the promoter was defined and the promoter region comprised between the 10th and 90th percentiles of the CAGE signal distribution was considered, as suggested in (Haberle *et al.*, 2015). Finally, TCs across all the analysed samples were aggregated, if supported by at least 5 tpm and closer than 100 bp, to form consensus clusters (CC) for downstream analyses. Having defined the final set of promoters (i.e., consensus clusters), the promoter density has been calculated as follows. First, by using the clusterdist algorithm of the ClusterScan tool (-d 10,000 parameter) (Volpe *et al.*, 2018), the zebrafish genome has been scanned grouping together all the consecutive promoters closer than 10 kb. Then, for each group of promoters (clusters), the promoter density has been calculated by dividing the number of promoters by the cluster length in kilobases. Finally, the rainfall plot representing the promoter density shown in **Figure 1E** has been obtained by using the karyoploteR R library (Gel and Serra, 2017).

### Processing of CAGE-seq data

CAGE-seq data set from PRJNA169500 (Nepal *et al.*, 2013) were mapped with Bowtie v1.1.2 (Langmead *et al.*, 2009) in default n-mode allowing 2 mismatches in the 27 bp seed region and reporting alignments of reads with up to 1 or 1000 valid alignments for unique (`-S -n 2 -l 27 -a --best -m 1`) and multimapping (`-S -n 2 -l 27 -a --best -m 1000`) respectively. The CAGEr package (Haberle *et al.*, 2015) was used for downstream processing of the data, i.e. CAGE Transcription Start Site (CTSS) calling and clustering into tag and consensus clusters (TCs and CCs).

### Processing of ChIP-seq and ATAC-seq data

Prior mapping ChIP-seq and ATAC-seq reads were trimmed from adapters and filtered for quality using TrimGalore (https://www.bioinformatics.babraham.ac.uk/projects/trim_galore) with automatic adaptor detection option and quality threshold of 20. Only reads where both mates survived the filtering steps were kept in case of paired end data: The data were mapped to a reference genome using segemehl aligner (v0.3.4) (Otto *et al.*, 2014) Allowing up to 2500 alignments per read and alignments accuracy of 90 %. (command line parameters: `-M 2500 -A 0.9`). Genome coverage was generated using STAR (Dobin *et al.*, 2013) with command line parameters: `--runMode inputAlignmentsFromBAM-outWigType bedGraph--outWigStrand Untranded--outWigNorm RPM`.

To compare the dynamics of the epigenetic features between the different gene set (**Figure 2A-B**) and datasets, the distribution of average normalised signal relative to the transcription start site for each set and feature was visualised on aggregation plots with the Bioconductor SeqPlots package (Stempor and Ahringer, 2016) providing the coverage file and genomic regions of features of interest as input with 20 bp bin size. For visualisation purposes a smoothing spline function was applied before plotting.

The transcription start sites (TSSs) were determined using the CAGE-seq at a developmental stage where there was sufficiently high signal to accurately determine the dominant TSSs for each gene in the set, or to minimize interference from the maternal contribution in the case of the constitutive gene set. For the miR-430, minor and major wave gene sets High, sphere and 30%-epiboly were used for dominant TSS detection, respectively. For both the constitutive and post-gastrulation sets Prim-5 stage was used.

The publicly available ChIP-seq and ATAC-seq data used in this study were obtained from the Sequence Read Archive (SRA) deposited under following BioProject accession: PRJNA434216 (H3K4me3 and H3K27me3 ChIP-seq); PRJNA473799 (H3K27ac ChIP-seq); PRJNA395463 (ATAC-seq); PRJNA156233 (Nanog ChIP-seq); PRJNA171706 (Pou5f1 and Sox2 ChIP-seq).

### Promoter architecture analyses

To compare core promoter motif composition between gene set (minor wave, major wave, constitutive and postulation) promoter sequences were extracted and centre-aligned to the dominant TSS.

Position weight matrices (PWM) for each core promoter set was calculated from the sequences using the Bioconductor “DiffLogo” package and motif sequence logos were visualised with the R “ggseqlogo” package.

Promoter architecture (sharp/broads), was determined for each gene in the above four datasets, using the 0.1-0.9 interquartile width (the length of the region in bp where 10%-90% of the CAGE signal is concentrated) of the promoter associated CAGE-tags consensus clusters. Promoters with interquartile width up to 10 bp were classified as sharp and with more than 10 bp as broad.

### Transcription factor binding motif analyses

Enrichment for clusters of TF binding sites was done using Cluster-Buster (Frith *et al.*, 2003) with the following parameters: cbust -r 10000 -m 5 -c 10 The optimal motif weights as abundances of the motifs (occurrences per kb) and the average distance between neighbouring motifs to use with Cluster-Buster was calculated using Cluster-Trainer (a supplementary tool for use with Cluster-Buster), ran with the following parameters:`ctrain -t 10 -r 10000`. As input sequence a 1.6Mb region (extracted form chr4 of the guided genome assembly of the Nanopore sequencing run, flanking 500 kb up- and downstream the miR-430 cluster. Position frequency matrices for Nanog (UN0383.1), Pou5f1::Sox2 (MA0142.1) and Sox2 (MA0143.3) where obtained from JASPAR(2020) database (http://jaspar.genereg.net)

### Generation of stable transgenic zebrafish reporter lines

The miR-430 promoter region fragment (~650 bp) was PCR amplified from zebrafish genomic DNA using the following forward: actcacgtgtggtaccCTTAGTCACCTCTGCCCACCAAG and reverse: catcgataagcggccgcTCCGTTCATAGTCTTTAGCAGCAAG primers (lower case letters correspond to the vector homology arms, upper case letters are specific to the miR-430 promoter regions) and cloned into into modified pJet:attB:mCherry vector (Roberts *et al.*, 2014) containing phiC31 attB site and mCherry reporter using In-Fusion HD Cloning Kit (TaKaRa/Clontech) according to the manufacture instructions. In brief the pJet vector was digested with Acc65I and NotI restriction enzymes. The purified linearized vector and miR-430 promoter fragments (in 1:3 molar ratio) were incubated with 1x In-Fusion HD cloning mix at 50°C for 15 min and 2 μl of the cloning reaction was transformed into the provided with the kit Stellar strain competent cells. Positive colonies were identified by PCR and the isolated plasmid verified by restriction digestion and sequencing.

Zebrafish reporter lines were generated by following the phiC31 integrase mediated targeted transgene integration protocol as described in (Hadzhiev *et al.*, 2016). One-cell stage docking line embryos were micro-injected with 1 nl solution containing the reporter plasmid (20 ng/μl) supplemented with phiC31*-nanos* 3’UTR mRNA (30 ng/μl). The efficiency of integration events was determined by frequency of lens colour change (from green to red) in the injected embryos at 72-96 hpf. If lens colour change is observed in more than 5 % of the embryos in an injection batch all the embryos from the batch are grown to adulthood, including those without lens colour change, because integration events may have happened with higher frequency in the PGCs due to *nanos* 3’UTR driven localisation of the phiC31 integrase mRNA occurring predominantly in the PGCs.

### Reporter expression analyses of miR-430 promoter stable transgenic lines by RT-PCR

Total RNA was extracted from ~50 embryos at the relevant developmental stage, using miRNeasy mini, (QIAGEN, 217004) following kit instructions. On-column DNAse treatment was performed using RNAse-free DNAse Set (QIAGEN, 79254). Reverse transcription was carid out using 1 μg of total RNA, 0.5 μg of random hexamers (Promega, C1181) and 200U M-MLV reverse transcriptase (Promega, M170A) in 25 μl reaction volume. PCR reactions were performed with GoTaq® Hot Start Master Mix (Promega, M5122) using 1 μl of the reverse transcription reaction as template in 20 μl reaction volume. Thirty amplification cycles were carried out in total and PCR reactions were analysed on agarose gel electrophoresis. The following primer pairs were used: miR430_Prom_FP: AGCAGACAACAAGATGCGTGTG with Gal4VP16_RP: ACTTCGGTTTTTCTTTGGAGCAC and γ-crystalline_Prom_FP GAAACTTCCACTCAGTCAGACTTGC with mCherry_RP: CACCTTGAAGCGCATGAACTC.

### Injections at single cell stage for ubiquitous delivery vs late-stage injection for mosaic treatments

For dCas9 and Cas9 targeting, a 1:1 ratio mix of two guide RNAs targeting the miR-430 repeats or one guide targeting the gol promoter was used.

Injection mixes for CRISPR mediated manipulations consisted of 650 ng/ul dCas9 protein (NEB, M0652) or Cas9 protein (NEB, M0646), and 200 ng/ul of guide RNA. For visualisation of CRISPRi knockdown reagents 400 ng/ul dCas9-GFP protein (Novateinbio, PR-137213G) and 200 ng/ul of guide RNA. These injection mixes were preincubated together at 37 °C for 5 minutes to facilitate complex formation, before supplementation with 0.1% phenol red and injection. Ubiquitous delivery of knock-down and labelling reagents was achieved by 1 nl injection of embryos at the one-cell stage. Mosaic knock-down was produced by 1 nl injection into a single cell at the 8-cell stage.

### Global transcription block with α-amanitin and triptolide treatment

Global transcription block was performed either by treatment with 2 μM triptolide from the one-cell stage (Sigma T3652) in E3 media, or by microinjection of 200 pg α-amanitin (Sigma A2263) at single-cell stage. As an injection control for α-amanitin, 10 % DMSO in water was used.

### PCNA staging

mApple:PCNA fusion protein (400 pg) was injected at the one-cell stage to enable ubiquitous labelling of all daughter nuclei for the selection of synchronous embryos. At the cell cycle of interest embryos were selected for ubiquitous bright, homogenous nuclear staining, indicative of nuclear-wide replication. These late interphase embryos were immediately snap frozen for RNA extraction or fixed with cold 4% PFA for in situ hybridisation.

### miR-430 MO labelling

Single-cell stage embryos were injected with 1 nl solutions containing Cy5-labelled miR-430 targeting morpholinos (Gene Tools LLC) (Hadziev *et al.*, 2019) at a concentration of 7 μM each in nuclease-free water and 0.1% phenol red.

### Ethynyl uridine (EU) labelling of nascent RNA

Single-cell stage zebrafish embryos were injected with 1 nl of 50 mM ethynyl-uridine (EU, Thermo Fisher, C10329) until fixation with 4% PFA at the stage of interest. Embryos were dehydrated and rehydrated using MeOH gradients then fluorescently labelled using the Click-iT™ RNA Alexa Fluor™ 488 Imaging Kit (Thermo Fisher, C10329), following the manufacturer’s protocol. Labelled embryos were used for subsequent protein RNA, or DNA detection.

### 3D DNA-FISH (miR-430 or klf17)

BACs DKEY-69C19 and CH1023-918E12 were used as templates for DNA-FISH probe production, these correspond to 214kb of the miR-430 region on chromosome 4 and 38kb of the *klf17* region on chromosome 2 respectively. The FISH Tag™ DNA Multicolor Kit (ThermoFisher) was used to label digested BAC fragments with Alexa dyes following kit instructions, with a 60 min digestion per μg of DKEY-69C19 and 10 min per μg of CH1023-918E12. 50-100 ng of probe was used for each sample in with hybridization buffer [50% formamide, 4x SSC, 100mM NaPO4 pH7.0, 0.1% Tween-20].Embryos were fixed at the developmental stage of interest using 4% PFA. Isolated animal caps were equilibrated with hybridization buffer, through a series of gradient washes. Embryonic DNA was then denatured at by 85°C for 10 minutes before immediate application of probe and incubation overnight at 37°C. A hybridization buffer: PBS-T gradient of washes was used to remove unbound probe from samples before counterstaining with DAPI. Samples were imaged on a Zeiss LSM 880 with FastAiryscan at maximum resolution, with a 63 ×1.40 numerical aperture objective lens. 20-50 optical sections (500 nm thickness) were acquired of each nuclei, and resulting images were processed using Zen Black software (Zeiss).

### RT-qPCR

Total RNA was extracted from >30 PCNA-synchronised 512-cell stage embryos, using miRNeasy mini, (QIAGEN,#217004) following kit instructions for optimised for small RNAs from animal cells. On-column DNAse treatment was performed using RNAse-free DNAse Set, (QIAGEN, 79254). Three independent experimental samples were collected for each manipulation condition and used for RT-qPCR analysis. 500ng of total RNA was used with M-MLV reverse transcriptase (Promega, M170A) to produce equal amounts of total RNA, using either miR-430/5S rRNA specific reverse primers or random hexamers (Promega, C118A) following manufacturer’s instructions. -RT controls were prepared alongside RT reactions with water replacing the reverse transcriptase. RT-qPCR reactions were performed using 10 ng of cDNA with 500 nM of each forward and reverse gene-specific primers and PowerUp SYBR Green MasterMix (Applied bio systems, A25742) on a QuantStudioV (Thermo Fisher Scientific) machine. Thermo Fisher Connect software was used for data acquisition and PRISM for statistical analysis. Ct values from independent biological repeats were evaluated by the Shapiro-Wilk test, which showed the data was normally distributed. The p-values were estimated by t-test. -RT controls gave Ct values either >7 cycles greater than +RT samples or equivalent to alpha-amanitin treated samples representing complete block of transcription.

### Guide RNA production

Double stranded DNA templates for sgRNA guides were produced through PCR annealing and extension of 100 ng common guide RNA backbone oligo and 100 ng target specific sgRNAR oligo (ordered from Merck) using Phusion Hot-start II DNA Polymerase (ThermoScientific, F549) with 35 cycles and annealing at 60 °C. Resultant DNA was gel extracted then 400 ng template used for in vitro transcription with the HiScribe RNA synthesis T7 kit (NEB, E2040) for 2 hours at 37 °C, then TurboDNase (Ambion, AM2238) treated for 15 minutes at 37 °C. Protein and non-incorporated nucleotides were removed using the Monarch RNA Clean up kit (NEB, T2040), before quantification by nanodrop.

### In situ probe production

Templates for in situ probes were produced by amplification from 100 ng of 1k cell stage cDNA using gene-specific primers with promoter overhangs and Phusion Hot-start II DNA Polymerase (ThermoScientific, F549). Resultant templates were gel extracted then 500 ng transcribed into RNA using T7 RNA polymerase (ThermoScientific, EP0111) and either dig- or FITC-labelling mix (Roche, 3935420 and 32874620 respectively). RNA probes were purified using Microspin G25 (GE Healthcare, 27-5325-01) columns then final RNA concentration determined by nanodrop. 30 ng of pre-exhausted probe in Hybe+ was used per 50 embryo sample, for RNA-ISH and-FISH.

### Live embryo imaging

The Zeiss Lightsheet Z1 was used to image live embryos mounted in 1% low melt agarose within size three glass capillaries, incubated at 28 °C in E3 media during imaging. Z-stacks of ~100–200 slices in 1 μm steps were acquired every 30–60 seconds for 1.5–2.5 h, with the ×20 objective.

### Whole mount antibody staining

Embryos were fixed at the 512-cell stage in 4% PFA then washed with PBS-0.3% TritonX-100 to permeabilize. 10% goat serum in PBST was used to block embryos before addition of pre-blocked Pol II S2P monoclonal antibody (Diagenode, C15200005) at 1:1000 dilution. PBS-0.1% Tween-20 washes removed excess primary antibody before application of pre-blocked Alexa 633 goat anti-mouse IgG (Life technologies, A21052) secondary antibody at 1:2000 dilution. Embryos were mounted in antifade mounting media with DAPI (Vectashield, H-1200, Vector Laboratories) and imaged with a Zeiss 880 confocal microscope with Fast Airyscan Module with a 63 ×1.40 numerical aperture objective lens and the resulting images were processed using Zen Black software (Zeiss).

### Whole mount In Situ Hybridisation

Embryos were fixed in 4% PFA at the developmental stage of interest, then washed with PBST before manual dechorionation and isolation of animal caps. Samples were dehydrated then rehydrated using a MeOH gradient before being equilibrated with hybridization buffer [50% formamide, 1.3x SSC, 5mM EDTA pH 8.0, 0.2% Tween-20]. Embryos were incubated with probe overnight at 58-65°C then unincorporated probe removed by a series of Hybe:SSCT and SSCT:PBST washes. Samples were then blocked in 10% goat serum in PBST before application of pre-blocked anti-dig-AP antibodies at a 1:2000 dilution in block. Excess antibody was removed with dig wash buffer (Roche, 11585762001) before staining with NBT and BCIP in 100mM NaCl, 100mM Tris.HCl pH9.5, 50mM MgCl2 & 0.1% Tween-20. Samples were imaged on a Zeiss AxioZoom.V16, with an ApoZ 1.5x/0.37 numerical aperture objective lens and the resulting images were processed using Zen Black software (Zeiss) and ImageJ.

## ACKNOWLEDGEMENTS

The presented work was funded by a Wellcome Trust Investigator Award (106955/Z/15/Z) to FM. This research was supported in part (SB) by the Intramural Research Program of the National Human Genome Research Institute (ZIAHG200386-06). We thank the MDS Technology Hub core at the University of Birmingham for flow cytometry and genomics support and the BMSU facility at the University of Birmingham for zebrafish maintenance. We thank Simon Branford and Jean-Baptiste Cazier for support and assistance with accessing the University of Birmingham BlueBEAR/CaStLeS high performance computing cluster. Ada Jimenez-Gonzalez for embryo injections during Covid19-associated facility restrictions, Piotr Balwierz for advice on gene copy number calculations and Michael Levine for coining the term ‘transcription organizer’ to describe our miR-430-associated observations.

## REFERENCES

Abe, K.I., Funaya, S., Tsukioka, D., Kawamura, M., Suzuki, Y., Suzuki, M.G., Schultz, R.M., and Aoki, F. (2018). Minor zygotic gene activation is essential for mouse preimplantation development. Proc Natl Acad Sci U S A 115, E6780–E6788. 10.1073/pnas.1804309115.

Alonge, M., Soyk, S., Ramakrishnan, S., Wang, X., Goodwin, S., Sedlazeck, F.J., Lippman, Z.B., and Schatz, M.C. (2019). RaGOO: fast and accurate reference-guided scaffolding of draft genomes. Genome Biol 20, 224. 10.1186/s13059-019-1829-6.

Amodeo, A.A., Jukam, D., Straight, A.F., and Skotheim, J.M. (2015). Histone titration against the genome sets the DNA-to-cytoplasm threshold for the Xenopus midblastula transition. Proc Natl Acad Sci U S A 112, E1086–1095. 10.1073/pnas.1413990112.

Ballarino, M., Pagano, F., Girardi, E., Morlando, M., Cacchiarelli, D., Marchioni, M., Proudfoot, N.J., and Bozzoni, I. (2009). Coupled RNA processing and transcription of intergenic primary microRNAs. Molecular and cellular biology 29, 5632–5638. 10.1128/mcb.00664-09.

Bartfai, R., Balduf, C., Hilton, T., Rathmann, Y, Hadzhiev, Y., Tora, L., Orban, L., and Muller, F. (2004). TBP2, a vertebrate-specific member of the TBP family, is required in embryonic development of zebrafish. Curr Biol 14, 593–598. 10.1016/j.cub.2004.03.034.

Blythe, S.A., and Wieschaus, E.F. (2015). Zygotic genome activation triggers the DNA replication checkpoint at the midblastula transition. Cell 160, 1169–1181. 10.1016/j.cell.2015.01.050.

Blythe, S.A., and Wieschaus, E.F. (2016). Establishment and maintenance of heritable chromatin structure during early Drosophila embryogenesis. Elife 5, e20148. 10.7554/eLife.20148.

Chan, S.H., Tang, Y., Miao, L., Darwich-Codore, H., Vejnar, C.E., Beaudoin, J.D., Musaev, D., Fernandez, J.P., Benitez, M.D.J., Bazzini, A.A., et al. (2019). Brd4 and P300 Confer Transcriptional Competency during Zygotic Genome Activation. Dev Cell 49, 867–881 e868. 10.1016/j.devcel.2019.05.037.

Chen, K., Johnston, J., Shao, W., Meier, S., Staber, C., and Zeitlinger, J. (2013). A global change in RNA polymerase II pausing during the Drosophila midblastula transition. Elife 2, e00861. 10.7554/eLife.00861.

Chen, Z., Omori, Y., Koren, S., Shirokiya, T., Kuroda, T., Miyamoto, A., Wada, H., Fujiyama, A., Toyoda, A., Zhang, S., et al. (2019). De novo assembly of the goldfish (Carassius auratus) genome and the evolution of genes after whole-genome duplication. Sci Adv 5, eaav0547. 10.1126/sciadv.aav0547.

Cisse, II, Izeddin, I., Causse, S.Z., Boudarene, L., Senecal, A., Muresan, L., Dugast-Darzacq, C., Hajj, B., Dahan, M., and Darzacq, X. (2013). Real-time dynamics of RNA polymerase II clustering in live human cells. Science 341, 664–667. 10.1126/science.1239053.

Collart, C., Allen, G.E., Bradshaw, C.R., Smith, J.C., and Zegerman, P. (2013). Titration of four replication factors is essential for the Xenopus laevis midblastula transition. Science 341, 893–896. 10.1126/science.1241530.

Conic, S., Desplancq, D., Ferrand, A., Fischer, V., Heyer, V., Reina San Martin, B., Pontabry, J., Oulad-Abdelghani, M., Babu, N.K., Wright, G.D., et al. (2018). Imaging of native transcription factors and histone phosphorylation at high resolution in live cells. J Cell Biol 217, 1537–1552. 10.1083/jcb.201709153.

Cook, P.R., and Marenduzzo, D. (2018). Transcription-driven genome organization: a model for chromosome structure and the regulation of gene expression tested through simulations. Nucleic Acids Res 46, 9895–9906. 10.1093/nar/gky763.

Creyghton, M.P., Cheng, A.W., Welstead, G.G., Kooistra, T., Carey, B.W., Steine, E.J., Hanna, J., Lodato, M.A., Frampton, G.M., Sharp, P.A., et al. (2010). Histone H3K27ac separates active from poised enhancers and predicts developmental state. Proc Natl Acad Sci U S A 107, 21931–21936. 10.1073/pnas.1016071107.

Davidson, A.E., Balciunas, D., Mohn, D., Shaffer, J., Hermanson, S., Sivasubbu, S., Cliff, M.P., Hackett, P.B., and Ekker, S.C. (2003). Efficient gene delivery and gene expression in zebrafish using the Sleeping Beauty transposon. Dev Biol 263, 191–202. 10.1016/j.ydbio.2003.07.013.

De Iaco, A., Planet, E., Coluccio, A., Verp, S., Duc, J., and Trono, D. (2017). DUX-family transcription factors regulate zygotic genome activation in placental mammals. Nature genetics 49, 941–945. 10.1038/ng.3858.

De Iaco, A., Verp, S., Offner, S., Grun, D., and Trono, D. (2020). DUX is a non-essential synchronizer of zygotic genome activation. Development 147. 10.1242/dev.177725.

Dobin, A., Davis, C.A., Schlesinger, F., Drenkow, J., Zaleski, C., Jha, S., Batut, P., Chaisson, M., and Gingeras, T.R. (2013). STAR: ultrafast universal RNA-seq aligner. Bioinformatics 29, 15–21. 10.1093/bioinformatics/bts635.

Ernst, J., Kheradpour, P., Mikkelsen, T.S., Shoresh, N., Ward, L.D., Epstein, C.B., Zhang, X., Wang, L., Issner, R., Coyne, M., et al. (2011). Mapping and analysis of chromatin state dynamics in nine human cell types. Nature 473, 43–49. 10.1038/nature09906.

Ferg, M., Sanges, R., Gehrig, J., Kiss, J., Bauer, M., Lovas, A., Szabo, M., Yang, L., Straehle, U., Pankratz, M.J., et al. (2007). The TATA-binding protein regulates maternal mRNA degradation and differential zygotic transcription in zebrafish. EMBO J 26, 3945–3956. 10.1038/sj.emboj.7601821.

Frith, M.C., Li, M.C., and Weng, Z. (2003). Cluster-Buster: Finding dense clusters of motifs in DNA sequences. Nucleic Acids Res 31, 3666–3668. 10.1093/nar/gkg540.

Gao, M., Veil, M., Rosenblatt, M., Gebhard, A., Hass, H., Buryanova, L., Yampolsky, L.Y, Grüning, B., Timmer, J., and Onichtchouk, D. (2020). Pluripotency factors select gene expression repertoire at Zygotic Genome Activation. bioRxiv, 2020.2002.2016.949362. 10.1101/2020.02.16.949362.

Gaskill, M.M., Gibson, T.J., Larson, E.D., and Harrison, M.M. (2021). GAF is essential for zygotic genome activation and chromatin accessibility in the early Drosophila embryo. Elife 10. 10.7554/eLife.66668.

Gehrig, J., Reischl, M., Kalmar, E., Ferg, M., Hadzhiev, Y., Zaucker, A., Song, C., Schindler, S., Liebel, U., and Muller, F. (2009). Automated high-throughput mapping of promoter-enhancer interactions in zebrafish embryos. Nature methods 6, 911–916. 10.1038/nmeth.1396.

Gel, B., and Serra, E. (2017). karyoploteR: an R/Bioconductor package to plot customizable genomes displaying arbitrary data. Bioinformatics 33, 3088–3090. 10.1093/bioinformatics/btx346.

Gentsch, G.E., Spruce, T., Owens, N.D.L., and Smith, J.C. (2019). Maternal pluripotency factors initiate extensive chromatin remodelling to predefine first response to inductive signals. Nat Commun 10, 4269. 10.1038/s41467-019-12263-w.

Giraldez, A.J., Mishima, Y., Rihel, J., Grocock, R.J., Van Dongen, S., Inoue, K., Enright, A.J., and Schier, A.F. (2006). Zebrafish MiR-430 promotes deadenylation and clearance of maternal mRNAs. Science 312, 75–79. 10.1126/science.1122689.

Green, M.R., and Sambrook, J. (2017). Isolation of High-Molecular-Weight DNA from Mammalian Tissues Using Proteinase K and Phenol. Cold Spring Harb Protoc 2017. 10.1101/pdb.prot093484.

Grow, E.J., Flynn, R.A., Chavez, S.L., Bayless, N.L., Wossidlo, M., Wesche, D.J., Martin, L., Ware, C.B., Blish, C.A., Chang, H.Y., et al. (2015). Intrinsic retroviral reactivation in human preimplantation embryos and pluripotent cells. Nature 522, 221–225. 10.1038/nature14308.

Guan, D., McCarthy, S.A., Wood, J., Howe, K., Wang, Y., and Durbin, R. (2020). Identifying and removing haplotypic duplication in primary genome assemblies. Bioinformatics 36, 2896–2898. 10.1093/bioinformatics/btaa025.

Haberle, V., Arnold, C.D., Pagani, M., Rath, M., Schernhuber, K., and Stark, A. (2019). Transcriptional cofactors display specificity for distinct types of core promoters. Nature 570, 122–126. 10.1038/s41586-019-1210-7.

Haberle, V., Forrest, A.R., Hayashizaki, Y., Carninci, P., and Lenhard, B. (2015). CAGEr: precise TSS data retrieval and high-resolution promoterome mining for integrative analyses. Nucleic Acids Res 43, e51. 10.1093/nar/gkv054.

Haberle, V., Li, N., Hadzhiev, Y., Plessy, C., Previti, C., Nepal, C., Gehrig, J., Dong, X., Akalin, A., Suzuki, A.M., et al. (2014). Two independent transcription initiation codes overlap on vertebrate core promoters. Nature 507, 381–385. 10.1038/nature12974.

Haberle, V., and Stark, A. (2018). Eukaryotic core promoters and the functional basis of transcription initiation. Nature reviews. Molecular cell biology 19, 621-637. 10.1038/s41580-018-0028-8.

Hadzhiev, Y., Miguel-Escalada, I., Balciunas, D., and Müller, F. (2016). Testing of Cis-Regulatory Elements by Targeted Transgene Integration in Zebrafish Using phiC31 Integrase. In Zebrafish: Methods and Protocols, K. Kawakami, E.E. Patton, and M. Orger, eds. (Springer New York), pp. 81–91. 10.1007/978-1-4939-3771-4_6.

Hadzhiev, Y., Qureshi, H.K., Wheatley, L., Cooper, L., Jasiulewicz, A., Van Nguyen, H., Wragg, J.W., Poovathumkadavil, D., Conic, S., Bajan, S., et al. (2019). A cell cycle-coordinated Polymerase II transcription compartment encompasses gene expression before global genome activation. Nat Commun 10, 691. 10.1038/s41467-019-08487-5.

Heintzman, N.D., Stuart, R.K., Hon, G., Fu, Y., Ching, C.W., Hawkins, R.D., Barrera, L.O., Van Calcar, S., Qu, C., Ching, K.A., et al. (2007). Distinct and predictive chromatin signatures of transcriptional promoters and enhancers in the human genome. Nature genetics 39, 311–318. 10.1038/ng1966.

Hendrickson, P.G., Dorais, J.A., Grow, E.J., Whiddon, J.L., Lim, J.W., Wike, C.L., Weaver, B.D., Pflueger, C., Emery, B.R., Wilcox, A.L., et al. (2017). Conserved roles of mouse DUX and human DUX4 in activating cleavage-stage genes and MERVL/ HERVL retrotransposons. Nature genetics 49, 925–934. 10.1038/ng.3844.

Heyn, P., Kircher, M., Dahl, A., Kelso, J., Tomancak, P., Kalinka, A.T., and Neugebauer, K.M. (2014). The earliest transcribed zygotic genes are short, newly evolved, and different across species. Cell Rep 6, 285–292. 10.1016/j.celrep.2013.12.030.

Hilbert, L., Sato, Y., Kuznetsova, K., Bianucci, T., Kimura, H., Julicher, F., Honigmann, A., Zaburdaev, V., and Vastenhouw, N.L. (2021). Transcription organizes euchromatin via microphase separation. Nat Commun 12, 1360. 10.1038/s41467-021-21589-3.

Hnisz, D., Shrinivas, K., Young, R.A., Chakraborty, A.K., and Sharp, P.A. (2017). A Phase Separation Model for Transcriptional Control. Cell 169, 13–23. 10.1016/j.cell.2017.02.007.

Howe, K., Schiffer, P.H., Zielinski, J., Wiehe, T., Laird, G.K., Marioni, J.C., Soylemez, O., Kondrashov, F., and Leptin, M. (2016). Structure and evolutionary history of a large family of NLR proteins in the zebrafish. Open Biol 6, 160009. 10.1098/rsob.160009.

Hug, C.B., Grimaldi, A.G., Kruse, K., and Vaquerizas, J.M. (2017). Chromatin Architecture Emerges during Zygotic Genome Activation Independent of Transcription. Cell 169, 216–228 e219. 10.1016/j.cell.2017.03.024.

Ichikawa, K., Tomioka, S., Suzuki, Y., Nakamura, R., Doi, K., Yoshimura, J., Kumagai, M., Inoue, Y, Uchida, Y, Irie, N., et al. (2017). Centromere evolution and CpG methylation during vertebrate speciation. Nat Commun 8, 1833. 10.1038/s41467-017-01982-7.

Joseph, S.R., Palfy, M., Hilbert, L., Kumar, M., Karschau, J., Zaburdaev, V., Shevchenko, A., and Vastenhouw, N.L. (2017). Competition between histone and transcription factor binding regulates the onset of transcription in zebrafish embryos. Elife 6, e23326. 10.7554/eLife.23326.

Kaaij, L.J.T., van der Weide, R.H., Ketting, R.F., and de Wit, E. (2018). Systemic Loss and Gain of Chromatin Architecture throughout Zebrafish Development. Cell Rep 24, 1–10 e14. 10.1016/j.celrep.2018.06.003.

Koren, S., Walenz, B.P., Berlin, K., Miller, J.R., Bergman, N.H., and Phillippy, A.M. (2017). Canu: scalable and accurate long-read assembly via adaptive k-mer weighting and repeat separation. Genome Res 27, 722–736. 10.1101/gr.215087.116.

Langmead, B., Trapnell, C., Pop, M., and Salzberg, S.L. (2009). Ultrafast and memory-efficient alignment of short DNA sequences to the human genome. Genome Biol 10, R25. 10.1186/gb-2009-10-3-r25.

Larson, M.H., Gilbert, L.A., Wang, X., Lim, W.A., Weissman, J.S., and Qi, L.S. (2013). CRISPR interference (CRISPRi) for sequence-specific control of gene expression. Nat Protoc 8, 2180–2196. 10.1038/nprot.2013.132.

Lee, M.T., Bonneau, A.R., and Giraldez, A.J. (2014). Zygotic genome activation during the maternal-to-zygotic transition. Annual review of cell and developmental biology 30, 581–613. 10.1146/annurev-cellbio-100913-013027.

Lee, M.T., Bonneau, A.R., Takacs, C.M., Bazzini, A.A., DiVito, K.R., Fleming, E.S., and Giraldez, A.J. (2013). Nanog, Pou5f1 and SoxB1 activate zygotic gene expression during the maternal-to-zygotic transition. Nature 503, 360–364. 10.1038/nature12632.

Leichsenring, M., Maes, J., Mossner, R., Driever, W., and Onichtchouk, D. (2013). Pou5f1 transcription factor controls zygotic gene activation in vertebrates. Science 341, 1005–1009. 10.1126/science.1242527.

Levine, M., Cattoglio, C., and Tjian, R. (2014). Looping back to leap forward: transcription enters a new era. Cell 157, 13–25. 10.1016/j.cell.2014.02.009.

Lindeman, L.C., Andersen, I.S., Reiner, A.H., Li, N., Aanes, H., Ostrup, O., Winata, C., Mathavan, S., Muller, F., Alestrom, P., and Collas, P. (2011). Prepatterning of developmental gene expression by modified histones before zygotic genome activation. Dev Cell 21, 993–1004. 10.1016/j.devcel.2011.10.008.

Lismer, A., Dumeaux, V., Lafleur, C., Lambrot, R., Brind’Amour, J., Lorincz, M.C., and Kimmins, S. (2021). Histone H3 lysine 4 trimethylation in sperm is transmitted to the embryo and associated with diet-induced phenotypes in the offspring. Dev Cell 56, 671–686 e676. 10.1016/j.devcel.2021.01.014.

Liu, G., Wang, W., Hu, S., Wang, X., and Zhang, Y (2018). Inherited DNA methylation primes the establishment of accessible chromatin during genome activation. Genome Res 28, 998–1007. 10.1101/gr.228833.117.

Lund, E., Liu, M., Hartley, R.S., Sheets, M.D., and Dahlberg, J.E. (2009). Deadenylation of maternal mRNAs mediated by miR-427 in Xenopus laevis embryos. RNA 15, 2351–2363. 10.1261/rna.1882009.

Macfarlan, T.S., Gifford, W.D., Driscoll, S., Lettieri, K., Rowe, H.M., Bonanomi, D., Firth, A., Singer, O., Trono, D., and Pfaff, S.L. (2012). Embryonic stem cell potency fluctuates with endogenous retrovirus activity. Nature 487, 57–63. 10.1038/nature11244.

Meier, M., Grant, J., Dowdle, A., Thomas, A., Gerton, J., Collas, P., O’Sullivan, J.M., and Horsfield, J.A. (2018). Cohesin facilitates zygotic genome activation in zebrafish. Development 145, dev156521. 10.1242/dev.156521.

Miao, L., Tang, Y, Bonneau, A.R., Chan, S.H., Kojima, M.L., Pownall, M.E., Vejnar, C.E., and Giraldez, A.J. (2020). Synergistic activity of Nanog, Pou5f3, and Sox19b establishes chromatin accessibility and developmental competence in a context-dependent manner. bioRxiv, 2020.2009.2001.278796. 10.1101/2020.09.01.278796.

Muller, F., Zaucker, A., and Tora, L. (2010). Developmental regulation of transcription initiation: more than just changing the actors. Curr Opin Genet Dev 20, 533–540. 10.1016/j.gde.2010.06.004.

Murphy, PJ., Wu, S.F., James, C.R., Wike, C.L., and Cairns, B.R. (2018). Placeholder Nucleosomes Underlie Germline-to-Embryo DNA Methylation Reprogramming. Cell 172, 993–1006 e1013. 10.1016/j.cell.2018.01.022.

Nepal, C., Hadzhiev, Y., Previti, C., Haberle, V., Li, N., Takahashi, H., Suzuki, A.M., Sheng, Y., Abdelhamid, R.F., Anand, S., et al. (2013). Dynamic regulation of the transcription initiation landscape at single nucleotide resolution during vertebrate embryogenesis. Genome Res 23, 1938–1950. 10.1101/gr.153692.112.

Otto, C., Stadler, P.F., and Hoffmann, S. (2014). Lacking alignments? The next-generation sequencing mapper segemehl revisited. Bioinformatics 30, 1837-1843. 10.1093/bioinformatics/btu146.

Palfy, M., Joseph, S.R., and Vastenhouw, N.L. (2017). The timing of zygotic genome activation. Curr Opin Genet Dev 43, 53–60. 10.1016/j.gde.2016.12.001.

Palfy, M., Schulze, G., Valen, E., and Vastenhouw, N.L. (2020). Chromatin accessibility established by Pou5f3, Sox19b and Nanog primes genes for activity during zebrafish genome activation. PLoS Genet 16, e1008546. 10.1371/journal.pgen.1008546.

Rada-Iglesias, A., Bajpai, R., Swigut, T., Brugmann, S.A., Flynn, R.A., and Wysocka, J. (2011). A unique chromatin signature uncovers early developmental enhancers in humans. Nature 470, 279–283. 10.1038/nature09692.

Roberts, J.A., Miguel-Escalada, I., Slovik, K.J., Walsh, K.T., Hadzhiev, Y., Sanges, R., Stupka, E., Marsh, E.K., Balciuniene, J., Balciunas, D., and Muller, F. (2014). Targeted transgene integration overcomes variability of position effects in zebrafish. Development 141, 715–724. 10.1242/dev.100347.

Sandelin, A., Carninci, P., Lenhard, B., Ponjavic, J., Hayashizaki, Y., and Hume, D.A. (2007). Mammalian RNA polymerase II core promoters: insights from genome-wide studies. Nat Rev Genet 8, 424–436. 10.1038/nrg2026.

Schulz, K.N., and Harrison, M.M. (2019). Mechanisms regulating zygotic genome activation. Nat Rev Genet 20, 221–234. 10.1038/s41576-018-0087-x.

Stempor, P., and Ahringer, J. (2016). SeqPlots - Interactive software for exploratory data analyses, pattern discovery and visualization in genomics. Wellcome Open Res 1, 14. 10.12688/wellcomeopenres.10004.1.

Tani, S., Kusakabe, R., Naruse, K., Sakamoto, H., and Inoue, K. (2010). Genomic organization and embryonic expression of miR-430 in medaka (Oryzias latipes): insights into the post-transcriptional gene regulation in early development. Gene 449, 41–49. 10.1016/j.gene.2009.09.005.

Treangen, T.J., and Salzberg, S.L. (2011). Repetitive DNA and next-generation sequencing: computational challenges and solutions. Nat Rev Genet 13, 36–46. 10.1038/nrg3117.

Turelli, P., Castro-Diaz, N., Marzetta, F., Kapopoulou, A., Raclot, C., Duc, J., Tieng, V., Quenneville, S., and Trono, D. (2014). Interplay of TRIM28 and DNA methylation in controlling human endogenous retroelements. Genome Res 24, 1260–1270. 10.1101/gr.172833.114.

Vastenhouw, N.L., Cao, W.X., and Lipshitz, H.D. (2019). The maternal-to-zygotic transition revisited. Development 146. 10.1242/dev.161471.

Vastenhouw, N.L., Zhang, Y., Woods, I.G., Imam, F., Regev, A., Liu, X.S., Rinn, J., and Schier, A.F. (2010). Chromatin signature of embryonic pluripotency is established during genome activation. Nature 464, 922–926. 10.1038/nature08866.

Veenstra, G.J., Destree, O.H., and Wolffe, A.P. (1999). Translation of maternal TATA-binding protein mRNA potentiates basal but not activated transcription in Xenopus embryos at the midblastula transition. Mol Cell Biol 19, 7972–7982. 10.1128/mcb.19.12.7972.

Veil, M., Yampolsky, L.Y., Gruning, B., and Onichtchouk, D. (2019). Pou5f3, SoxB1, and Nanog remodel chromatin on high nucleosome affinity regions at zygotic genome activation. Genome Res 29, 383–395. 10.1101/gr.240572.118.

Volpe, M., Miralto, M., Gustincich, S., and Sanges, R. (2018). ClusterScan: simple and generalistic identification of genomic clusters. Bioinformatics 34, 3921 – 3923. 10.1093/bioinformatics/bty486.

Westerfield, M., and ZFIN. (2000). The zebrafish book: a guide for the laboratory use of zebrafish Danio (Brachydanio) rerio, 4th Edition (ZFIN).

White, R.J., Collins, J.E., Sealy, I.M., Wali, N., Dooley, C.M., Digby, Z., Stemple, D.L., Murphy, D.N., Billis, K., Hourlier, T., et al. (2017). A high-resolution mRNA expression time course of embryonic development in zebrafish. Elife 6, e30860. 10.7554/eLife.30860.

Wike, C.L., Guo, Y., Tan, M., Nakamura, R., Shaw, D.K., Diaz, N., Whittaker-Tademy, A.F., Durand, N.C., Aiden, E.L., Vaquerizas, J.M., et al. (2021). Chromatin architecture transitions from zebrafish sperm through early embryogenesis. Genome Res 31, 981–994. 10.1101/gr.269860.120.

Wragg, J., and Muller, F. (2016). Transcriptional Regulation During Zygotic Genome Activation in Zebrafish and Other Anamniote Embryos. Adv Genet 95, 161–194. 10.1016/bs.adgen.2016.05.001.

Wu, S.F., Zhang, H., and Cairns, B.R. (2011). Genes for embryo development are packaged in blocks of multivalent chromatin in zebrafish sperm. Genome Res 21, 578–589. 10.1101/gr.113167.110.

Xu, C., Fan, Z.P., Muller, P., Fogley, R., DiBiase, A., Trompouki, E., Unternaehrer, J., Xiong, F., Torregroza, I., Evans, T., et al. (2012). Nanog-like regulates endoderm formation through the Mxtx2-Nodal pathway. Dev Cell 22, 625–638. 10.1016/j.devcel.2012.01.003.

Yamazaki, Y., Eguchi, S., Miyamura, H., Hayashi, J., Fukuda, J., Ozeki, H., Yoshimura, T., Fujita, Y., and Tsuchiya, A. (1989). Replacement of myocardium with a Dacron prosthesis for complications of acute myocardial infarction. J Cardiovasc Surg (Torino) 30, 277–280.

Yang, H., Luan, Y., Liu, T., Lee, H.J., Fang, L., Wang, Y, Wang, X., Zhang, B., Jin, Q., Ang, K.C., et al. (2020). A map of cis-regulatory elements and 3D genome structures in zebrafish. Nature 588, 337–343. 10.1038/s41586-020-2962-9.

Zabidi, M.A., Arnold, C.D., Schernhuber, K., Pagani, M., Rath, M., Frank, O., and Stark, A. (2015). Enhancer-core-promoter specificity separates developmental and housekeeping gene regulation. Nature 518, 556–559. 10.1038/nature13994.

Zhang, B., Wu, X., Zhang, W., Shen, W., Sun, Q., Liu, K., Zhang, Y., Wang, Q., Li, Y., Meng, A., and Xie, W. (2018). Widespread Enhancer Dememorization and Promoter Priming during Parental-to-Zygotic Transition. Mol Cell 72, 673–686 e676. 10.1016/j.molcel.2018.10.017.

Zhu, I., Song, W., Ovcharenko, I., and Landsman, D. (2021). A model of active transcription hubs that unifies the roles of active promoters and enhancers. Nucleic Acids Res 49, 4493–4505. 10.1093/nar/gkab235.

Zhu, W., Xu, X., Wang, X., and Liu, J. (2019). Reprogramming histone modification patterns to coordinate gene expression in early zebrafish embryos. BMC Genomics 20, 248. 10.1186/s12864-019-5611-7.

