## Supplementary Figures for "The miR-430 locus with extreme promoter density is a transcription body organizer, which facilitates long range regulation in zygotic genome activation"

1.Institute of Cancer and Genomics Sciences, Birmingham Centre for Genome Biology, College of Medical and Dental Sciences, University of Birmingham, Birmingham, UK.
2. Area of Neuroscience, Scuola Internazionale Superiore di Studi Avanzati (SISSA), Trieste, Italy.
3. Central RNA Laboratory, Istituto Italiano di Tecnologia (IIT), Genova, Italy.
4. South China Sea Institute of Oceanology, Chinese Academy of Sciences, Guangzhou, China.
5.Translational and Functional Genomics Branch, National Human Genome Research Institute, National Institutes of Health, Bethesda MD, USA.
*These authors contributed equally

### SUPPLEMENTARY FIGURES

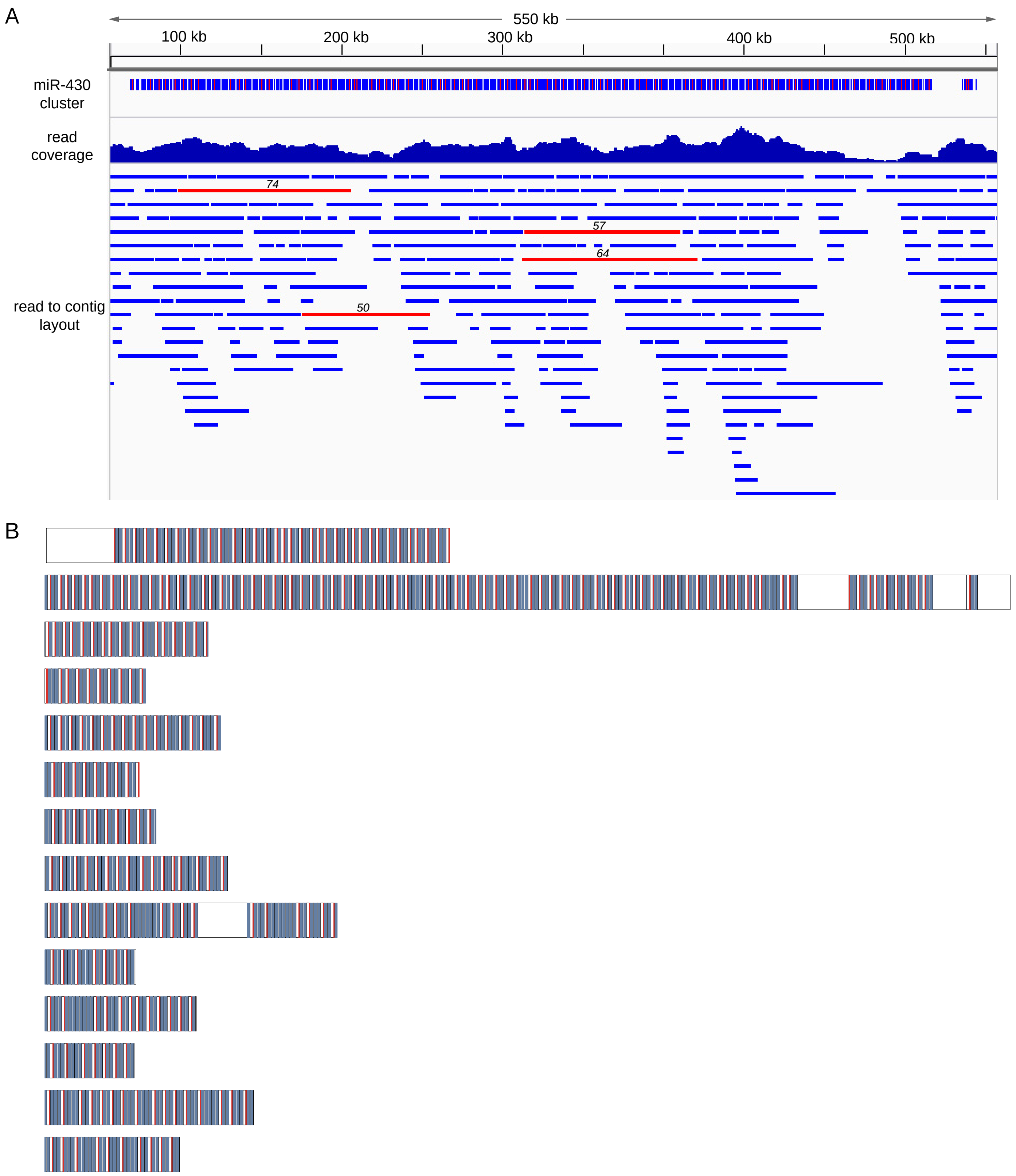


**Supplementary Figure 1.** ***De novo*** **assembly of the miR-430 cluster from long read sequencing data.**
**A,** Genome Browser screenshots of the miR-430 gene cluster on contig2 assembled from Nanopore long reads. The three tracks (top to bottom) show the structure of the cluster (promoters in red, precursor triplets in blue), long read coverage and the read to contig assembly layout, respectively. Raw reads in the read to contig layout track with 50 or more promoters are in red with the promoter number indicated on top. **B,** Schematic representation of miR-430 containing contigs from the de novo assembly of the PacBio sequencing. Red boxes represent the miR-430 core promoter, blue the precursor triplet.

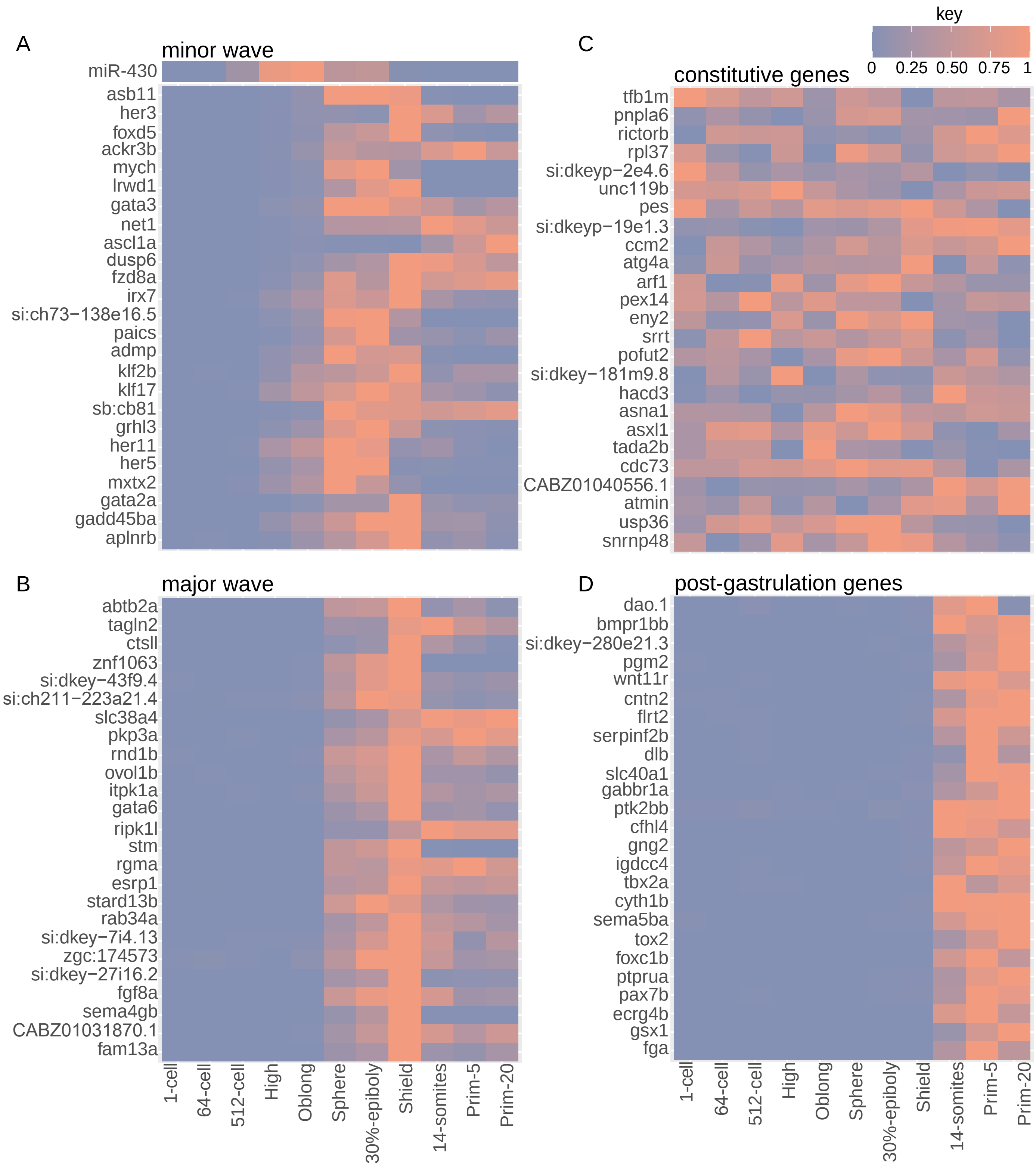


**Supplementary Figure 2 Expression dynamics of miR-430 and the 4 gene set used in this study as determined from CAGE-seq.**

**A**, miR-430 and minor wave. **B,** major wave. **C,** constitutive genes. **D,** post-gastrulation genes. For visualisation purposes the CAGE tag per million (tpm) values were scaled from 0 to 1.

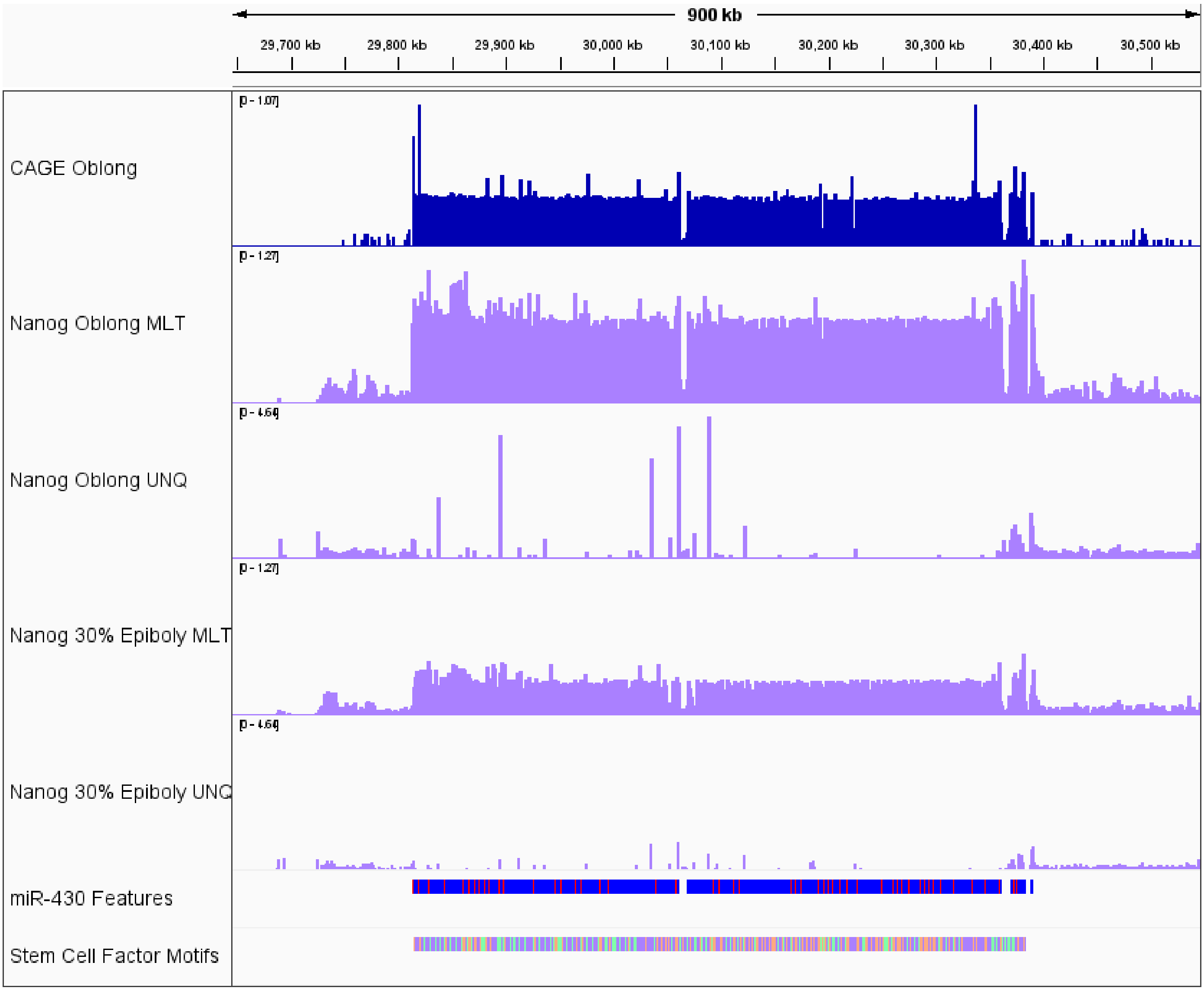


**Supplementary Figure 3. Overview of CAGE-seq, Nanog ChIP-seq and predicted stem sell factor binding sites distribution on miR-430 cluster.**

**A,** browser screenshot showing ChIP-seq signal coverage of the pluripotency factor Nanog (purple) for each stage unique (UNQ) and multi-mapped (MLT) read coverage is shown. CAGE-seq signal (top track, blue) marks the position of the TSS of each promoter. Bottom tracks show the miR-430 features and the position of pluripotency factor binding sites, predicted by Cluster-Buster.

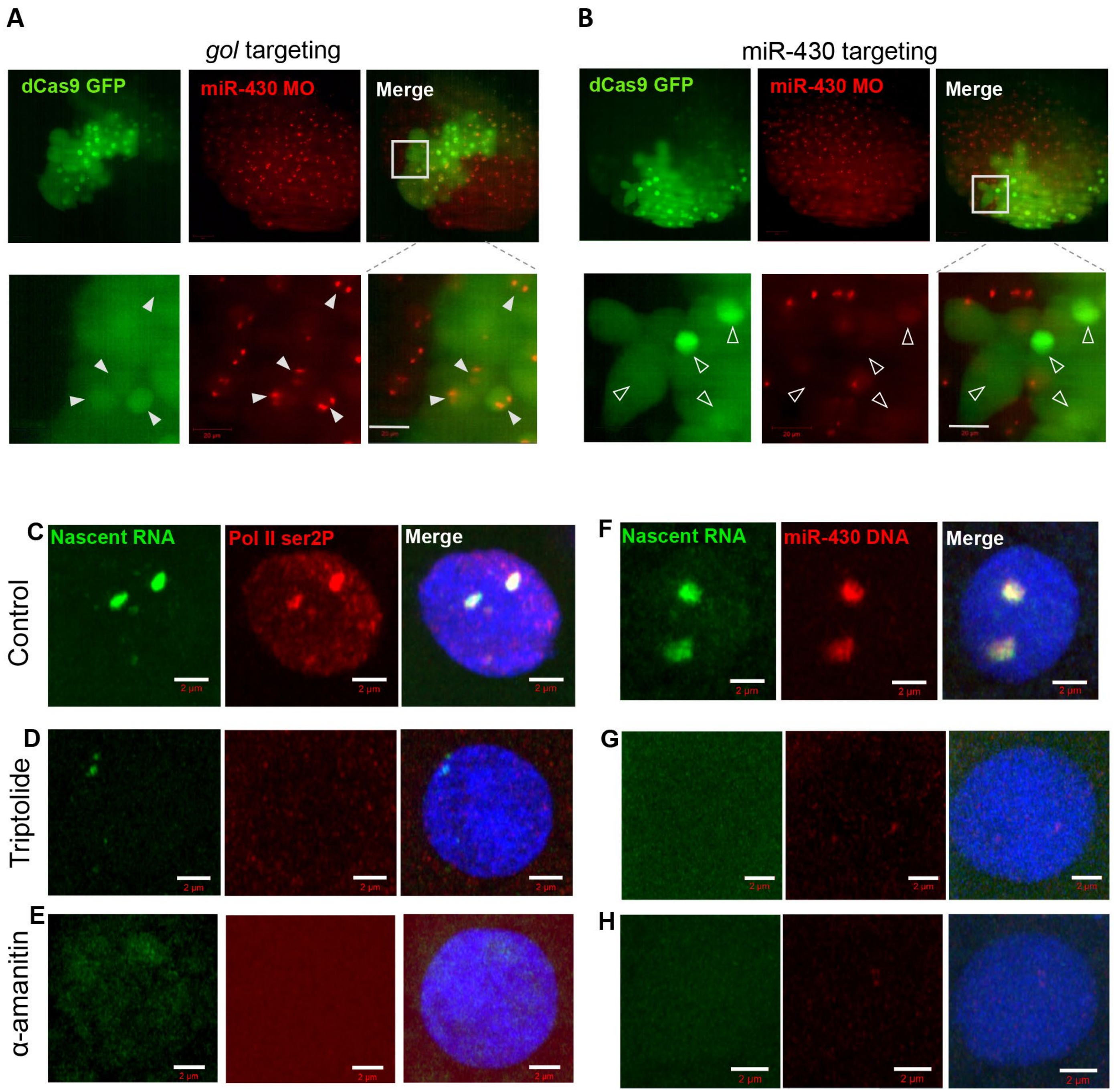


**Supplementary Figure 4. CRISPRi manipulation targeting the miR-430 cluster and global transcription block treatments causes loss of the transcription body.**

**A-B**, Live cell images of miR-430 MO (red) labelled nascent miR-430 RNA in 512 cell stage embryos following either gol splice junction (A) or miR-430 promoter (B) mosaic targeting with dCas9-GFP (green). Grey boxes indicate enlarged regions shown below. gol targeted: 10 embryos, miR-430 targeted: 8 embryos. Scale bars represent 20μm. **C-E**, EU labelling of nascent RNA (green), Pol II Ser2P (red) immunostaining and DAPI (blue) in 512-cell stage embryos. **C,** Injection control embryos, 60 nuclei from 6 embryos. **D,** Triptolide treated embryos, 55 nuclei from 7 embryos. **E,** α-amanitin injected embryos, 53 nuclei from 6 embryos. **F-H,** EU labelling of nascent RNA (green) combined with FISH for miR-430 DNA (red) and DAPI (blue) at the 512-cell stage. **F,** Injection control embryos, 31 nuclei from 5 embryos. **G,** Triptolide treated embryos, 38 nuclei from 6 embryos. **H,** α-amanitin injected embryos, 33 nuclei from 7 embryos. C-H, Scale bars represent 2μm.

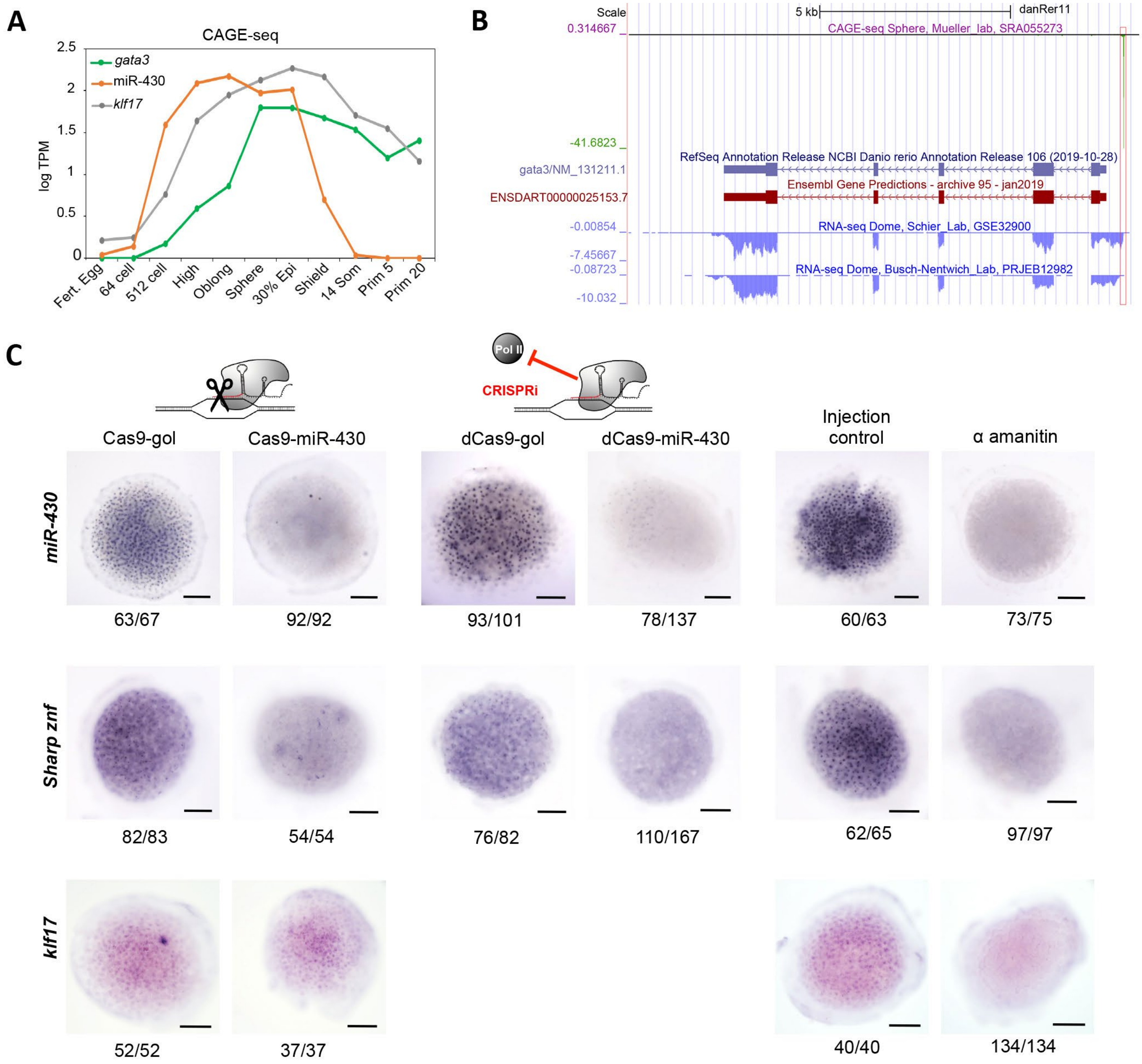


**Supplementary Figure 5. miR-430 manipulation affects early expression of neighbouring sharp *znf* genes.**
**A,** Promoter activity during early development by CAGE-seq, for miR-430, *gata3* and *klf17*. **B**, CAGE-seq and RNA-seq data suggest inaccurate annotation of the *gata3* transcript start by ENSEMBL and RefSeq. The determined by CAGE-seq and RNA-seq transcription start site (marked by red box) is ~500 bp upstream of the annotated transcript start. **C,** Whole animal cap top down views of expression patterns by chromogenic ISH of mir430, sharp promoter *znfs* and *klf17* at the 512-cell stage following various manipulations. Numbers under images indicate frequency of staining patterns in stained embryos. Scale bar = 100µm.
